# CXCL10/CXCR3 Signaling Induces Neural Senescence and Cognitive Impairments

**DOI:** 10.1101/2024.10.16.618653

**Authors:** Monika Patel, Sakesh Kumar, Aditya Singh, Prem N Yadav

**Affiliations:** Divison of Neuroscience & Ageing Biology, CSIR-Central Drug Research Institute, Lucknow-226031, India; Academy of Scientific and Innovative Research (AcSIR), Ghaziabad-201002, India

**Keywords:** CXCR3, Neural senescence, Cognitive impairments, Neurodegeneration, CXCL10, Aging

## Abstract

Chemokine receptors belong to the G-protein-coupled receptors family and multiple lines of emerging evidence suggest that several chemokines are elevated in aging associated with central nervous system disorders. Increased level of CXCL10 in central nervous system is reported in several neurodegenerative diseases, including Multiple sclerosis, Alzheimer’s disease, and Virus-associated dementia. We also observed significantly increased expression of CXCL10 and CXCR3 in the prefrontal cortex and hippocampus of aged C57BL/6J mice (12- and 18-month-old mice). This leads us to hypothesize that CXCL10, being a component of SASPs, may aggravate/perpetuate the brain aging process and, finally, neurodegenerative diseases. To test this hypothesis, we administered CXCL10 (intracerebroventricular: ICV, 0.5 pg/ hrs, 28 days) in 8-month-old C57BL/6J mice. We observed increased expression of senescent marker proteins p16^INK4a^, p21^Cip1^, p53 and decreased expression of pRB in the prefrontal cortex, which was blocked by CXCR3-specific antagonist AMG487. Furthermore, chronic infusion of CXCL10 induced learning and memory deficits in Y-maze, social recognition, contextual freeze tests and c-FOS expression in the prefrontal cortex. To further determine the specificity of CXCL10/CXCR3 signaling, we treated the primary cortical neuron (Days *in vitro*: DIV 7-8) with CXCL10 and found increased senescence in CXCR3 dependent fashion. Using RFP-EGFP-LC3-tagged transgenic mice, we also showed CXCL10/CXCR3 signaling attenuates autophagic flux in primary cortical neurons. Lastly, using a c-FOS-iRFP reporter, we observed that increased CXCL10/CXCR3 signaling impairs glutamatergic signaling in primary cortical neurons. These results support the hypothesis that increased CXCL10/CXCR3 facilitates brain aging and could be targeted for the management of aging-associated CNS disorders.

## Introduction

Advances in medical science and the health care system have increased the average life expectancy of the world population from 66.5 to 73.4 years in the last two decades (https://www.worldometers.info/demographics/life-expectancy). On the other hand, aging is one of the major risk factors for several chronic diseases, including neurodegenerative diseases, cardiovascular diseases, and type-2 diabetes (Madabhushi et al., 2014; Niccoli and Partridge, 2012; Yan et al., 2021).

Aging is actually due to a breakdown in the control mechanisms that are required in the complex performance of each tissue and organ. During the last decade, cellular senescence has emerged as a fundamental aging mechanism that contributes to diseases of late life, including neurodegenerative diseases such as Alzheimer’s disease (AD) and Parkinson’s disease (PD) (van Deursen, 2014). Multiple lines of evidence have demonstrated that senescent cells acquire “senescence-associated secretory phenotype (SASP)” and thereby recruit inflammatory cells, induce cell death, induce fibrosis, and inhibit resident stem cell functions (Baker et al., 2016; Jeon et al., 2017). Therefore, the development of SASP inhibition strategies and molecules for the clearance of senescent cells (senolytics) has gained significant traction recently (Bussian et al., 2018; Chang et al., 2016; Gill et al., 2022; Laberge et al., 2015).

Chronic low-grade sterile neuroinflammation in diseases such as AD and PD is considered one of the underlying mechanisms of neurodegenerative diseases (Bussian et al., 2018; Chang et al., 2016; Laberge et al., 2015; Ransohoff, 2016; Yong et al., 2024). Initiating factors for neuroinflammation are usually disease-specific, but there is significant convergence in the mechanisms responsible for the sensing, transduction, and amplification processes that ultimately lead to a disease. Chemokines, a subset of cytokines, have been shown to play significant roles in brain aging through various mechanisms, including inflammation, cellular senescence, and synaptic plasticity (Benveniste, 1992; Ramesh et al., 2013).

Increased levels of chemokines (CXCL10, and cytokines and their receptors, including IL-1α, CXCR2, CCR3, CCR5, and TGF-β, have been reported in AD (Cartier et al., 2005). Among all chemokines, CCL2, CX3CL1, and CXCL10 are most abundantly expressed in the brain (Cardona et al., 2008). Among these chemokines, CXCL10 is most widely implicated in neuroinflammatory and neurodegenerative conditions (Bradburn et al., 2018; Roberts et al., 2015; Spittau et al., 2012). CXC10 is a secreted peptide of 10 kDa, which was first discovered as an early response gene induced after interferon-γ treatment in cells and thus named an “interferon-inducible peptide 10 (IP-10)” (Luster and Ravetch, 1987; Luster et al., 1985). The increased CXCL10 expression is also shown to play an important role in viral infection-induced dementia and neuronal apoptosis (Kolb et al., 1999; Sui et al., 2004). Furthermore, CXCL10 is also shown to induce neuronal apoptosis *in vitro* via activation of intracellular Ca^++^ and caspase pathway (Sui Y et al.,2006). Given that the cognate receptor of CXCL10, CXCR3, primarily couples to the Gαi family of proteins, this study didn’t clarify if the CXCL10 effect was direct or indirect on intracellular Ca^++^ increase. Furthermore, systemic injection of viral mimetic leads to a robust increase in CXCL10/CXCR3 signaling, followed by cerebral hyperexcitability and altered synaptic plasticity (Petrisko et al., 2020). However, it is not known if the subtle but sustained increase in CXCL10 in the brain can induce aging pathways and associated behavioral outcomes.

In this study, we also observed significantly increased expression of CXCL10 and CXCR3 in 12- and 18-month-old mice in comparison to adult mice (2-month-old mice). This leads us to hypothesize that CXCL10, being a component of SASPs, may aggravate/perpetuate the brain aging process and, consequently, cognitive impairments. To test this hypothesis, we treated the primary cortical neuron days *in vitro* (DIV-7-8) with CXCL10 and found increased senescence in a CXCR3-dependent fashion. Also, CXCL10 decreased the autophagic flux, and increased lipid droplets and β-gal staining in primary cortical neurons. Detailed *in vivo* studies in mice revealed that chronic brain infusion of CXCl10 in normal C57BL/6J mice leads to increased expression of senescent marker proteins p16^INK4a^, p21^Cip1^, and p53 and decreased expression of pRB, which were blocked by CXCR3 specific antagonist AMG487. Furthermore, we observed that chronic infusion of CXCL10 induced learning and memory deficits in Y-maze, social recognition, and contextual freeze tests.

## Material and Methods

### Animal

The experiments and procedures were conducted *in vivo* in accordance with the guidelines provided by the Institutional Animal Ethics Committee (IAEC) of CSIR-Central Drug Research Institute, Lucknow, India. Male C57BL/6J mice aged 2M, 8M, 12M, and 18M, weighing between 22-25 g, were utilized for this study.

These mice were kept in a controlled environment with a 12-hour light/dark cycle, consistent humidity (50 ± 5%), and a constant temperature of 22-25°C. They were provided with *ad libitum* access to food and water.

### Drugs

CXCL10, an endogenous CXCR3 agonist (R&D, 266-IP/CF), was initially dissolved in 1XPBS at a concentration of 20μg/ml. It was then diluted to (2 ng/ml) with normal ACSF, and 200 µl of this was loaded into the ALZET Osmotic Pump, sufficient to release at the rate of .25 µl/hrs for 28 days. The CXCR3 selective antagonist AMG487 (Tocris, 4487) was initially dissolved in DMSO at a concentration of 10 mM diluted to a working concentration of 1.3 µM and loaded into the Alzet pumps along with CXCL10. Control animals were injected with 0.9% sodium chloride (vehicle).

### Osmotic pump ICV infusion of CXCL10 or AMG487 using mini ALZET osmotic pumps

We implanted a mini osmotic pump (ALZET Osmotic Pumps, model no. 2004) along with an ICV cannula in mice to deliver either CXCL10 (R&D, 266-IP/CF) alone or in a combination of AMG487 (Tocris, 4487). To do this, we used C57BL/6J mice and inserted a 30-gauge intra-cerebral ventricular cannula (from Plastic One, Roanoke, USA) into the right ventricle of the mice’s brain. The mice were then infused with CXCL10 (2 ng/ml) and CXL10+AMG487 (1.3 µM). The animals were anesthetized using ketamine and xylazine (90 mg/kg and 10 mg/kg, respectively) prior to the procedure. We marked the right ventricle (located at XY coordinates: −0.5mm anterior, posterior, and −1.1mm medial, lateral) using the bregma as the reference point (Nascimento et al., 2016). The cannula and guide cannula were cut according to the right ventricle coordinate (−2.2 mm depth). The cannula was then firmly implanted in the skull with the use of an ALZET osmotic pump and dental cement.

After the dental cement had hardened, the remaining skin adhered with tissue adhesive (Vet Bond, 3M Science, USA). Following the surgical procedures, the mice were allowed to rest for 3–4 hours in a warm cage under a heating lamp (with a maintained temperature of around 30-32 °C). The animals were given ibuprofen (0.04 mg/ml in drinking water) and gentamycin (1mg/kg, i.p.) for the first three days to alleviate pain and prevent infection. Subsequently, the mice were allowed to acclimate to the implanted cannula for one week.

### Cytokines Array

We profiled the levels of cytokines and chemokines in various age mice PFC (2M,12M, and 18M) using the (Procarta PlexTM, PPX-11-MXH6CM4) following the manufacturer’s instructions. Briefly, a 100-microgram protein lysate or standard was incubated with 50 µL of the capture beads for 2 hours at room temperature on a shaker. Thereafter, we washed the plate three times and added 25 µL of detection antibody mix (1X), again incubated with shaking at room temperature for 30 minutes. We washed the plate three times and then added 50 µL of streptavidin-PE and further incubated it with shaking at room temperature for 30 minutes. Finally, the plate was washed three times, and the beads were resuspended. We added 120 µL of reading buffer, sealed the plate again, and shook it at room temperature for 5 minutes. Finally, we acquired data on the Luminex xMAP Intelligence System (Thermofisher Scientific), and the level of each cytokine or chemokine was calculated using a 4-parameter logistic curve drawn with a standard used with each cytokine or chemokine.

### Enzyme-linked immunosorbent Assays (ELISA)

CXCL10 ELISA kit (R&D Systems, DY466-05) was used to measure the levels of mouse CXCL10 following the manufacturer’s instructions. Firstly, we homogenized mouse cortical tissue lysates in lysis buffer (150 mM NaCl, 50 mM Tris, 1%Triton X-100) with protease inhibitors. Then, we centrifuged the homogenized mixture at 10000g for 20 minutes at 4°C and collected the supernatants, which were stored at - 80°C until further use. Next, we coated a 96-well microplate with the diluted capture antibody according to the kit instructions and sealed the plate for overnight incubation at room temperature. The following day, the microplate was washed three times with wash buffer (0.05% Tween 20 in PBS pH 7.2) and blocked with blocking buffer (1% BSA in PBS pH 7.2) for one hour at room temperature.

Thereafter, we incubated the plate with 100 µg of tissue lysates for 2 hours at room temperature. We then washed the microplate twice with wash buffer and added the detection antibody for 2 hours. We repeated the washing steps and incubated the plate with conjugated streptavidin-HRP for 30 minutes at room temperature, followed by washing the plate and adding substrate solution for 20 minutes at room temperature. Lastly, we added a stop solution to each well and read the plate with a microplate reader at 450 nm. The data was analyzed by Flex Station 3, a molecular device.

### Immunohistochemistry and confocal imaging

C57BL/6J mice were perfused with a solution of 4% paraformaldehyde through their hearts. After removal, their brains were fixed overnight in 4% paraformaldehyde and then dehydrated in 30% sucrose for 24 hours. Thin coronal slices of 20μm were obtained using a cryostat (FSE, ThermoScientific). To stain the slices, they were washed four times in PBS and subjected to antigen retrieval in 10mM sodium citrate buffer (pH-6.3), followed by permeabilization using 0.5% Triton X-100 in 1X PBS for one hour at room temperature. After that, the slices were blocked with a buffer consisting of 3% bovine serum albumin, 3% horse serum, and 0.3% Triton X-100 in PBS for two hours at room temperature. Finally, the slices were incubated overnight at 4°C with rabbit anti-CXCR3 (Abcam, ab71864) with dilution 1:500, anti-mouse CXCL10 (Abcam, ab9938) 1:500, anti-rabbit c-Fos (Abcam, ab7963) 1:500 and anti-mouse NeuN (Cell Signaling Technology, 94403) 1:500 and anti-rabbit NeuN (Abcam, ab128886),1:500. After overnight incubation, the brain sections were washed three times with PBST (0.1% Triton-X 100 in 1X PBS). Following that, the sections were incubated for one hour at room temperature with the corresponding secondary antibodies that were conjugated with Alexa fluor 594 or 488 (1:500) and Hoechst 33258 (2 µg/ml in a blocking buffer. Thereafter, the brain sections were mounted using a Vectashield antifade mounting media onto glass slides coated with Poly-L-lysine after three rounds of washing. Utilizing an Olympus BX61-FV1200-MPE microscope with a 40X oil (1.3 NA) objective and a scan speed of 8.0μs/pixel, (Olympus, Shinjuku, Tokyo, Japan) fluorescent pictures were acquired using the XYZ acquisition mode at a step size of 1μm. Pixel intensity quantification was carried out using ImageJ software (www.rcb.info.nih.gov/ij) for CXCR3 (Abcam, ab71864), CXCL10 (Abcam, ab9938), c-Fos (Abcam, ab7963), and NeuN staining in PFC and Hippocampus. After normalization with the pixel intensity of NeuN in each region, results were shown as fold change.

### Immunocytochemistry and confocal imaging for primary cortical neuron

Primary cortical neurons were prepared from 0–1-day-old pups and the cells were fixed with 2% PFA for 10 minutes at 12 days *in vitro* (DIV). To stain the culture, we washed them four times in PBS and permeabilized them using 0.3% Triton X-100 in 1X PBS for an hour at room temperature. Next, we blocked the primary cortical neuron using a buffer containing 3% bovine serum albumin, 3% horse serum, and 0.3% Triton X-100 in PBS for two hours at room temperature. Finally, we incubated primary culture overnight at 4°C using anti-rabbit-CXCR3 (Abcam, ab71864) at a dilution of 1:500, anti-mouse GFP at a dilution of 1:500, anti-mouse NeuN (Cell Signaling Technology, 94403) at a dilution of 1:500, and anti-rabbit p16^INK4a^ (Invitrogen, PAZ-20379) at a dilution of 1:500. After overnight incubation, primary cortical neurons were washed three times with PBST (0.1% Triton X-100 in 1X PBS) followed by incubation with appropriate secondary antibodies conjugated Alexa Fluor-594 or −488 (1:1000 dilutions) and Hoechst 33258 (2 µg/ml) in blocking buffer for 1 hours at room temperature. Fluorescent images were collected with XYZ acquisition mode at 1μm step size using the Olympus BX61-FV1200-MPE microscope, which used a 40X oil (1.3 NA) objective at 8.0 μs/pixel scan speed (Olympus, Shinjuku, Tokyo, Japan). To quantify the pixel intensity for CXCR3, p16^INK4a^, GFP, and NeuN staining in primary cortical neurons, pixel intensity quantification was performed using ImageJ software (www.rcb.info.nih.gov/ij). The results were presented as fold changes after normalization with the pixel intensity of NeuN and GFP. To stain primary neurons, the PFA fixed neurons were first blocked using a blocking buffer containing 3% BSA, 3% horse serum, and 0.3% Triton (in 1X PBS) for two hours at room temperature. Then, they were incubated with the following primary antibodies: anti-Rabbit p16^INK4a^ (Invitrogen, PAZ-20379) 1:500, anti-mouse NeuN (Millipore, A60-MAB377) 1:200, anti-rabbit NeuN (Abcam, ab128886) 1:1000 and anti-mouse MAP2 (Sigma, M9942)1:1000 for 16 hours at 4°C. After that, species-specific Alexa Fluor-488, and 594 (1:1000; Molecular Probes) tagged secondary antibodies were applied for two hours at room temperature. Finally, fluorescence signals were captured under a fluorescence microscope equipped with high sensitivity.

### Western blotting

Western blotting was performed to determine the expression of various proteins, as previously described (Dogra et al., 2016). Frozen brain tissues were homogenized in tissue lysis buffer (25 mM Tris-HCl, pH 7.6, 150 mM NaCl, 0.5% NP-40, 1% sodium deoxycholate, and 0.1% SDS) and 40–50 µg of protein samples were subjected to 8–12% SDS-PAGE and transferred to a polyvinyl difluoride (Pall, 0.2-micron PVDF) membrane. The membrane was then blocked for two hours at room temperature using a blocking buffer (5% non-fat dry milk and 3% bovine serum albumin in PBST). Specific primary antibodies anti-rabbit-p16^INK4a^ (Invitrogen, PAZ-20379 1:1000), anti-mouse p21^Cip1^ (Invitrogen, AHZ0422 1:1000) and anti-rabbit LC3 (Sigma, L7543 1:1000), anti-rabbit pRB (CST 9307, 1:1000) and anti-mouse p53 (Invitrogen MA5-12571, 1:1000) were incubated overnight at 4°C, followed by horseradish peroxidase (HRP)-conjugated secondary antibodies (1:4000) at room temperature for 2 hours. The blots were developed using an enhanced chemiluminescence substrate, and images were acquired using Amersham ImageQuantTM 800. Densitometric analysis was performed using ImageJ software.

### Synaptic plasticity study in mice with c-Fos expression

The adeno-associated viral (AAV) vector plasmid pOTTC476-pAAVc-Fos-eYFP (Addgene Plasmid #47907) was packaged into AAV9 serotype using pXX6-80 and pXR9 (kind gift from Dr. Aravind Asokan, UNC-Chapel Hill, USA) in HEK293 as described earlier (Dogra et al., 2016). Briefly, pAAV-c-Fos-eYFP virions were purified using a discontinuous iodixanol gradient (Sigma-Aldrich) and concentrated to 1 X 10^12^ genome copy/ml, and 0.5 µl of these viral particles were delivered bilaterally by stereotaxic injection in the mPFC of a mouse brain, AP=+1.96mm, DV=-2.28mm, and ML=±0.062mm using a motorized stereotaxic system (Stoelting, USA). Stereotaxic surgeries were performed using a mixture of ketamine and xylazine (90 mg/kg and 10 mg/ kg, respectively) as anesthetic agents and Meloxicam (1mg/ kg, three consecutive days, s.c.) as postoperative analgesia.

Further, the region specificity of the viral expression was verified in each mouse using immunohistochemistry. Mice expressing inadequate viral expression were excluded from the analysis.

### Primary cortical neuronal culture and shRNA-mediated knockdown of CXCR3

Primary cortical neurons were cultured according to the protocol described in our previous study (Dogra et al., 2016). In brief, cerebral cortices were isolated from 0–1-day-old mouse pups and digested using 0.1% papain (Sigma-Aldrich, P4762) for 20 minutes at 37°C. Once incubation was complete, enzyme activity was stopped by adding complete neurobasal media (supplemented with 2% B27 supplement and Glutamax 1X, Invitrogen). Next, the digested tissue samples were triturated 10-15 times using a Pasteur pipette to obtain a single-cell suspension. The triturated samples were allowed to sit for 2–3 minutes to enable the heavier cellular debris to settle down. The supernatant was then transferred to a fresh tube and spun at 200 g for 5 minutes to obtain the cellular pellet. The cell pellet was suspended in fresh complete neurobasal medium and plated at a density of 0.5 x 10^6^ cells per well in a 12-well plate. The neurons were subjected to various ligands of the CXCR3 receptor after 10–11 days *in vitro* (DIV). For shRNA-mediated knockdown of CXCR3, lentiviral particles containing CXCR3-shRNA (TL500382 V, Origine) were added at 5 days *in vitro* (DIV) in primary cortical neurons for 72 hours and for c-Fos expression we were added pOTTC475-pAAV-c-Fos-iRFP (Addgene plasmid-#47906) at DIV 8. All drug treatments were performed after 72 hours of lentiviral transduction, and neuronal cell lysates were prepared as described earlier (Dogra et al., 2016).

### Reverse transcription (RT) and quantitative PCR

Total RNA was extracted from the primary cortical neuron using the trizol/chloroform method. Thereafter total RNA was treated with DNase at 37°C for 10 minutes to remove genomic DNA, then reverse transcribed into cDNA (iScript TM cDNA Synthesis Kit #1708891). This cDNA was then used for real-time PCR detection using the turbo-GFP or CXCR3-specific primers and condition for PCR was 95°C for 10 minutes, followed by 40 cycles of 95°C for 15s and 56°C for 30s. The housekeeping gene “turbo-GFP” forward 5 TACTACAGCTCCGTGG 3’ and reverse primers 5’ TTCTTCACCGGCATCTGCAT 3’ was used as a reference gene for the normalization of gene expression “CXCR3” using primer forward 5’ GGATCTATTTCCGGTGAATTCGCCACCATCAA 3’ and reverse 5’ CTAGAACTAGTCTCGAGGAATTCCTAGAGGCCAGAATAACTAGCC 3’. The 2-ΔΔCT method (i.e. delta-delta-ct analysis) was used for relative quantification (Valasek and Repa, 2005).

## Behavioral Analysis

### Y-Maze

We investigated the effect of the CXCL10/CXCR3 axis on spontaneous short-term novelty-based spatial memory using the Y-maze test (Lauritzen et al., 2016). The dimensions of the maze’s arms were 10 cm in height, 7 cm in breadth, and 35 cm in length **(Figure 2.4)**. In brief, the animals were placed in the start arm and allowed freely to investigate the start arm as well as the familiar arm for five minutes during the exposure phase, but the third arm was blocked in this phase.

After the exposure phase, animals were returned to the home cage and were kept there for 10 min. During the test phase, the animals were returned to the beginning arm and allowed five minutes to explore both arms (novel and familiar arm). Each arm’s duration was recorded using the Any-maze software version 7. The floor was covered with corn bedding, which was changed every time a trial was conducted.

The spatial cues were preserved during the test phases. The following formula was used to calculate the discrimination index for each mouse:

Discrimination index = (time spent in the novel arm - time spent in the familiar arm)/total exploration time for both arms.

### Social Recognition

As stated before, a cubical box with dimensions of 40 × 40 × 40 cm was used to test social recognition (Leger et al., 2013) **(Figure 2.5)**. In brief, the C57BL/6J mice were habituated to the arena for 10 minutes without any objects in the testing arena. Following a 24-hour interval, the animals participated in a 10-minute test session with both a novel (C57BL/6J) and a familiar (C57BL/6J) mouse, as well as a 10-minute familiarization session. The exploring duration of both sessions (in seconds) was recorded using the Any-maze software version 7. The discrimination index for the novel object was calculated with the following equation:

Discrimination index= time of novel mice exploration - the time of familiar mice exploration/ total exploration time for both mice.

### Contextual Fear Conditioning

This behavioral test was conducted in an isolation cubicle (Habitats H10-24TA, Coulbourn Instruments) that was shielded from light and sound. A precise animal shocker (H13-15, Coulbourn Instruments) was used to provide the 0.7-mA scrambled foot shock. A typical sound card was used to create the 5000-Hz, 80-to 85-dB sinus tone sound stimuli. A HiFi amplifier (PR530A, Pyramid) was used to amplify the sound and transmit it through speakers mounted in the walls of chambers (contexts A). Context A was 17 cm by 18 cm by 32 cm in size, with two clear walls and stainless-steel grid floors (H10-11M-TC, Coulbourn Instruments). A personal computer and Freeze-frame software (Actimetrics Software) were used to control the devices. The tested animal’s movements were captured by a digital video camera fixed to the cubicle ceiling, and Freeze View software (Actimetrics Software) was used to calculate the percentage of freezing.

The virtual threshold for freezing was set at 10 in each experiment. Fear Conditioning: The protocol was created in accordance with research that had already been published (Cain et al., 2002). Day 1 (acquisition): C57BL/6J mice (Test and Vehicle) were placed in context A and are typically allowed 2 min to freely roam about the conditioning chamber. Conditional fear was induced by presenting three pairings in the form of white light, and an auditory cue was offered for 30 seconds, which co-terminated with a foot shock for 2 seconds. Two minutes of stimulus-free periods preceded, separated, and followed the pairings. After the last foot shock, the mice are left alone in their chambers for ninety seconds. Day 2 (recall): 24 hours later, mice were placed in the same arena with the exact same context to measure context-dependent recall of fear and hence freezing. Freezing was analyzed for 5 minutes with freeze frame software (Coulbourn, habit isolation cubicle).

### Locomotor Activity

The purpose of this test was to quantify the locomotor activity by Panlab Infrared (IR) Actimeter (IR Actimeter system, Panlab) as described earlier (Seibenhener et al.,2015) with slight modifications. Briefly, mice were put in an open-field arena (45 × 45 × 40 cm) and allowed to explore the entire arena for 60 minutes, to investigate the entire space. Following the 60-minutes testing period, the animals were taken out of the arena. The whole area was cleaned with water and a smell-free detergent solution to remove the olfactory cues of previous animals after every session.

## Results

### 1. Brain CXCL10 levels directly correlate with aging and cognitive impairments

Aging is widely linked to inflammaging, a protracted and heightened state of systemic inflammation (Baylis et al., 2013; Franceschi and Campisi, 2014). Multiple lines of evidence have unequivocally shown that low-grade sterile and chronic inflammation leads to cognitive decline and the risk of dementia (Kipnis et al., 2004; Ziv et al., 2006). Similarly, studies with aged heterochronic parabiosis mouse models exhibiting elevated plasma levels of the chemokine CCL11 (eotaxin) resulted in decreased adult neurogenesis and impaired spatial learning and memory (Villeda et al., 2011). Furthermore, epigenetic upregulation of systemic CXCL10 expression in the prefrontal cortex was also reported to be associated with cognitive impairments in older adults and patients of Alzheimer’s disease (AD), respectively (Bradburn et al., 2018). However, the causative relationship between CXCL10 expression and cognitive impairments, and mechanisms is still unclear. Therefore, in this study first, using a protein array, we investigated the levels of a panel of nine chemokines and cytokines (Eotaxin, RANTES, CXCL10, IL-13, IL17A, IL-6, TNF-α, IL-1β, and INFγ) in young (2-Month-old) and older (12- and 18-month-old) C57BL/6J mice prefrontal cortex (PFC) region, a brain region critically involved in cognition.

We observed a significant increase of CXCL10 in the PFC of mice in an age-dependent manner **(Figure 1A)**. The levels of eotaxin (CCL11) also found to be robustly increased in PFC in an age-dependent manner that has been reported earlier, albeit only in the plasma of older mice and AD patient’s immunohistochemistry and found a significant increase in older mice (12 and 18 months old) in comparison to young mice **(Figure 1B, C, D, and Figure S1)**.

**Figure 1:**
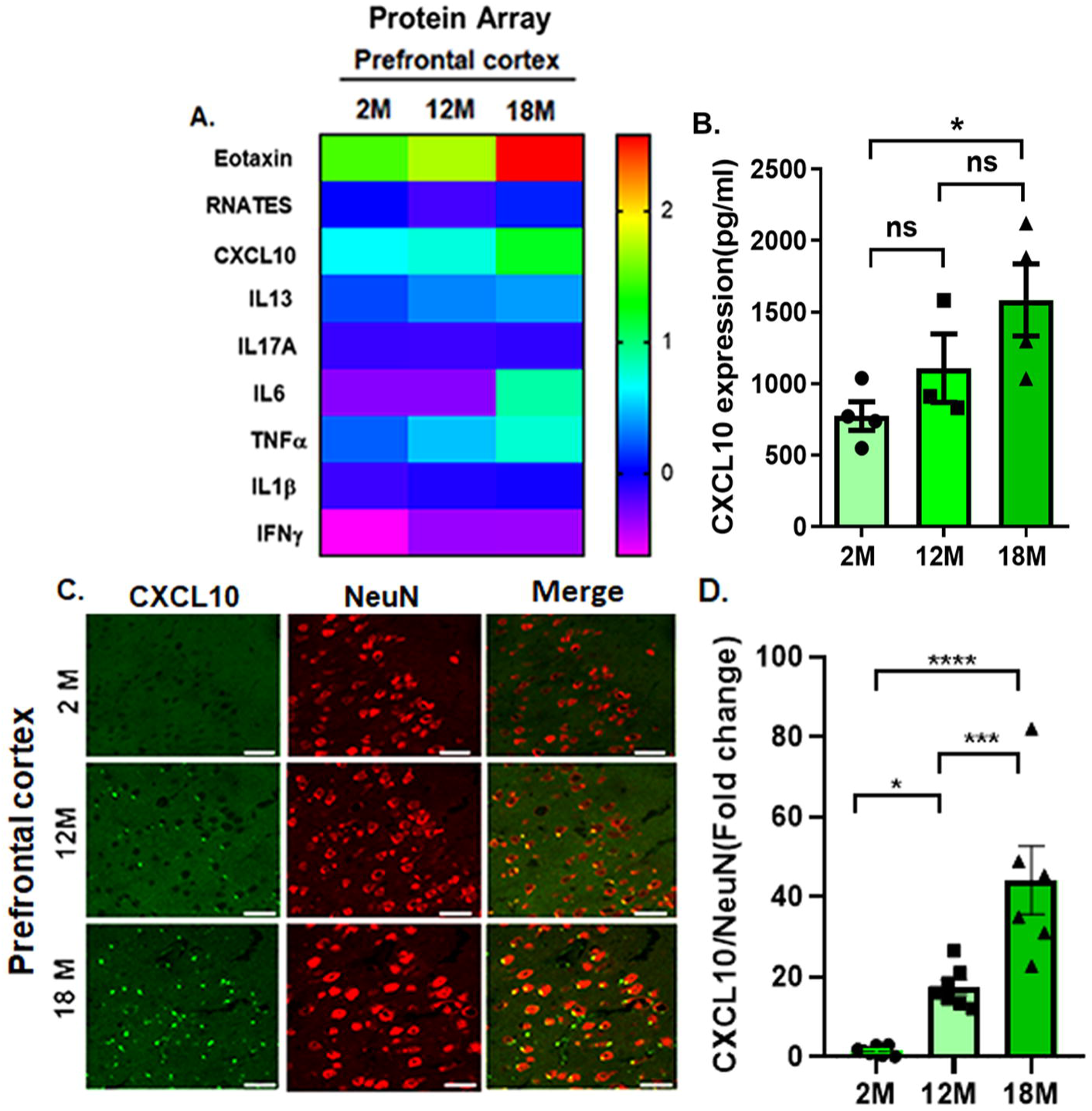
CXCL10 expression increased in an age-dependent manner in the PFC. (A) Heat map showing various cytokines and chemokines expression in different age groups of mice, red color showing high, and blue color showing low expression in PFC. A robust increase in PFC levels of eotaxin, followed by CXCL10, was observed. (B) A significant increase in the level of CXCL10 in PFC was also observed by ELISA, unpaired t-test, t=2.986, *p<0.05, 2M vs. 18M, *p<0.05, 12M vs. 18M n=3–4 mice/group. (C) Representative confocal images (40X magnification) of PFC showing increased CXCL10 expression in 2M, 12M, and 18M old C57BL/6J mice, CXCL10 (green) co-localizing with the neuronal marker NeuN (red), scale bar, 20µm. (D) Bar graph showing quantification of relative fluorescent intensity of CXCL10, normalized to NeuN intensity. Data were plotted as the mean ±SEM, one-way ANOVA followed by Tukey’s multiple comparisons test, F _(2, 21)_ = 11.42, ****p<0.001, 2M vs. 18M, ***p<0.01, 12M vs. 18M, n= 6-7 mice/ group.

Interestingly, we also found increased expression of CXCR3, the cognate receptor of CXCL10, in both PFC and CA3 of the hippocampus **(Figure S2)**. Given that both PFC and hippocampus are very crucial for optimal cognition, we also measured cognitive functions using Y-maze, social recognition, and contextual freezing in young and older age mice. We observed significant impairment in spatial learning and memory Y-maze **(Figure 2B)**, social recognition **(Figure 2C)**, and contextual freezing response **(Figure 2D)** in 18-month-old mice in comparison to young mice. These cognitive parameters in 12-month-old mice were also in decreasing trends, but not statistically significant in comparison to young mice. To further gain mechanistic insight underlying these cognitive impairments, we determined the levels of senescence marker proteins (p16^INK4a^, p21^Cip1^) and autophagy marker protein microtubule-associated protein 1A/1B-light chain 3 (LC3-βII) in PFC. Interestingly, we observed a significant increase in p16^INK4a^ and p21^Cip1^ and a decrease in LC3-βII in 18-month-older mice in comparison to 2-month-young mice **(Figure 2E-J)**. These results indicate the direct relationship between levels of CXCL10 in PFC and brain senescence and cognitive impairments in older mice.

**Figure 2:**
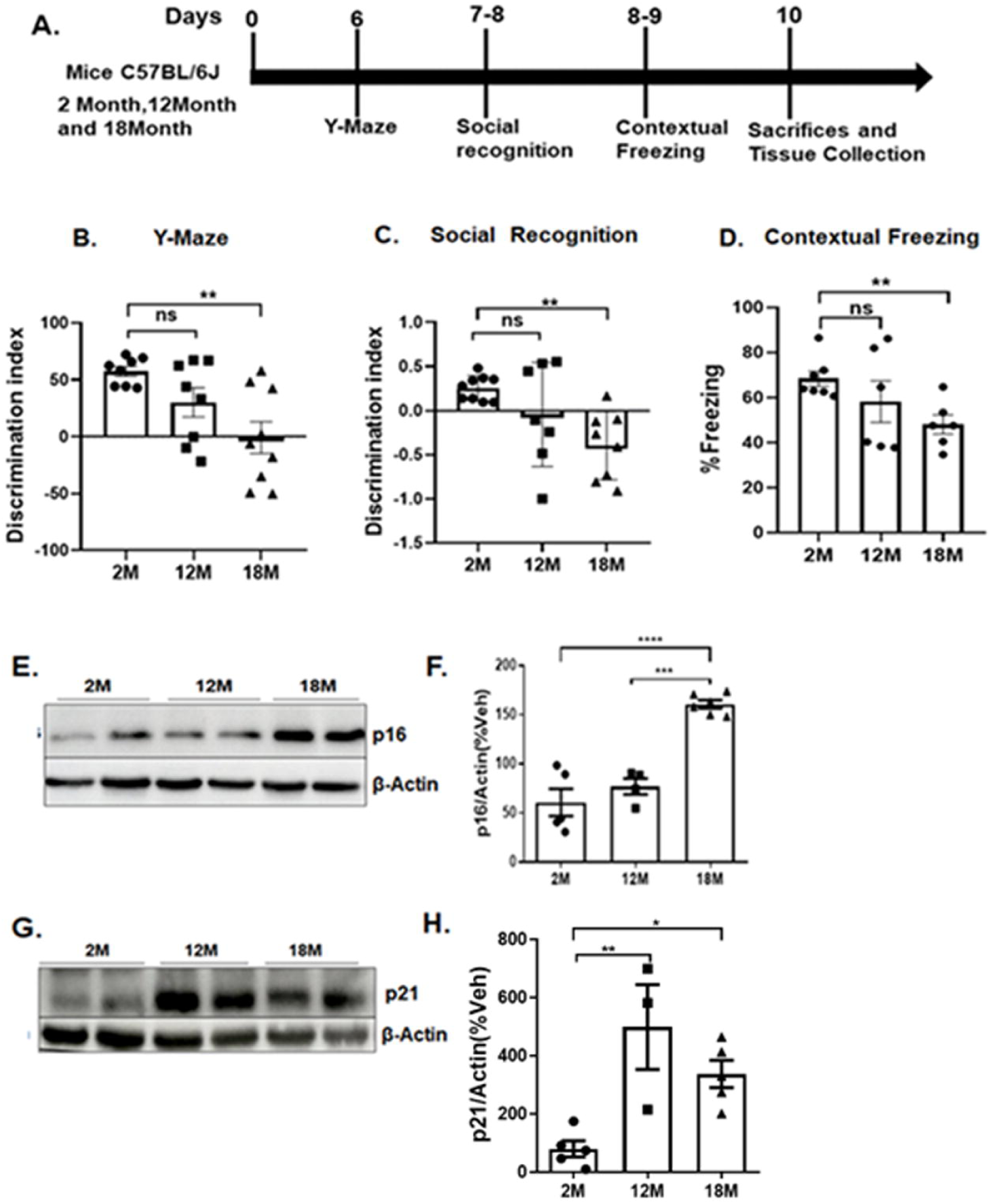
Cognitive impairments in aged mice are associated with increased senescence. (A) Diagram showing the timeline of the drug treatment regimen and behavioral assay in the C57BL/6J mice. (B) Bar graph showing the discrimination index between the novel and familiar arm Y-maze test in the different age groups of mice. Data were plotted as a mean ±SEM, one-way ANOVA followed by Tukey’s multiple comparisons test, F _(2, 20)_ = 7.830, **P<0.01, 2M vs. 18M, n=8-9 Mice/group. (C) Bar graph showing the discrimination index between the novel and familiar mice in the social recognition test. Data were plotted as a mean ±SEM one-way ANOVA followed by Tukey’s multiple comparisons test, F _(2, 26)_ =6.320, **p<0.01, 2M vs. 18M, n=7-9 mice/group. (D) Bar graph showing the % freezing due to contextual fear in the different age groups of mice. Data were plotted as mean ± SEM, unpaired t-test, **p<0.01, t=4.228, 2M vs. 18M, 6-7 mice/group. (E) Representative immunoblots show increased expression of p16^INK4a^ in an age-dependent manner in 2M,12M, and 18M old mice. (F) Bar graph showing quantification of p16^INK4a^ normalized to β-Actin, presented as percent of vehicles (%Veh). Data were plotted as a mean ± SEM, one-way ANOVA, followed by Tukey’s multiple comparisons test, F _(2,12)_ =36.49, ****p<0.0001, 2M vs. 18M, ***p<0.001, 12M vs.18M, n=4-6 mice/group. (G) Representative immunoblots showing increased expression of p21^Cip1^ in 12M and 18M-old mice in comparison to 2M-old mice (H) Bar graph showing quantification of p21^Cip1^ normalized to β-Actin and presented as percent vehicle (%Veh). Data were plotted as a mean ± SEM, one-way ANOVA, followed by Tukey’s multiple comparisons test, F _(2,10)_ =7.191 **p< 0.01, 2M vs.12M, *p<0.05, 2M vs.18M, n=3-5 mice/group.

### 2. CXCL10 induces brain senescence and cognitive impairments via CXCR3

Since the increased level of CXCL10 in PFC and hippocampus correlated with cognitive impairments, we sought to determine the specificity and mechanisms of CXCL10 action *in vivo*. To do this, we infused very modest CXCL10 (ICV, 0.5pg/ hrs, 28 days) either alone or in combination with CXCR3 selective antagonist AMG487 (ICV, 0.5pg/ hrs, 28 days) and then measured the learning and memory followed by PFC tissue analysis of senescence marker proteins. We observed significant impairment of spatial learning and memory in the Y-maze **(Figure 3B)** and social recognition memory **(Figure 3C)** and contextual freezing **(Figure 3D)** in CXCL10 infused mice, which were blocked by CXCR3 antagonist AMG487 except for the social recognition. Suggesting the majority of the neurobehavioral effect of CXCL10 is through its cognate receptor CXCR3 signaling. To further rule out the chronic effect of CXCL10 infusion on general exploratory behaviors of mice that can be a confounding factor in the learning and memory tests interpretations, we performed the open field (a novel arena for these mice) locomotor tests. As shown in **Figure 3E**, chronic infusion of CXCL10 in the brain doesn’t change the exploratory behaviors measured by the total horizontal distance traveled during 60 minutes. Furthermore, we observed that chronic treatment with CXCL10 significantly increased the PFC expression of senescence marker proteins p16^INK4a^, p21^Cip1^, p53, and pRB **(Figure 3F-M)**, which were blocked by CXCR3 antagonist AMG487. These observations suggest that increased levels of CXCL10 in the brain reinforce senescence mechanisms through its cognate receptor CXCR3 and consequently cause cognitive impairments.

**Figure 3:**
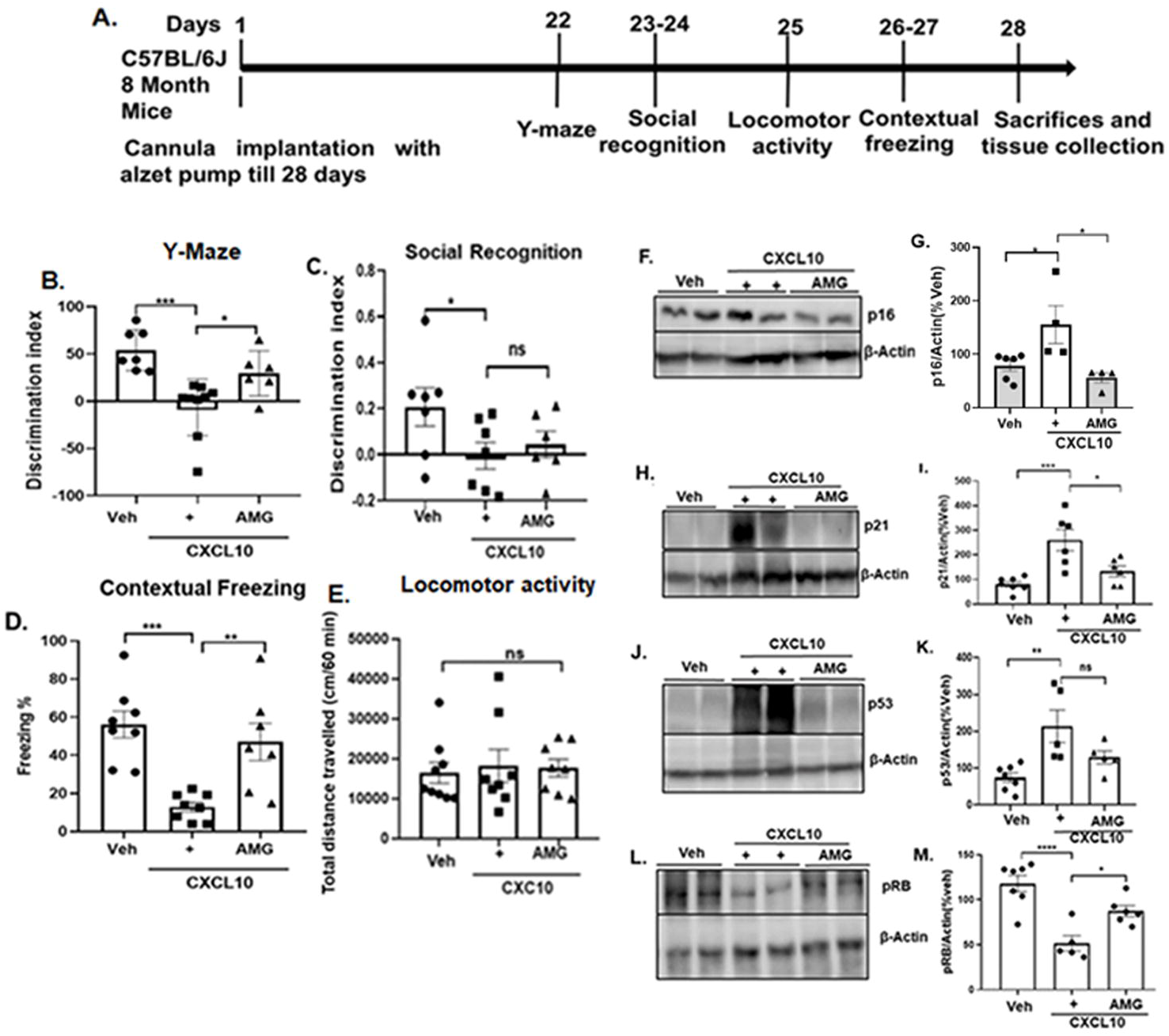
CXCR3/CXCL10 signaling impairs cognition and induced senescence in adult C57BL/6J mice. (A) Schematics showing the timeline of the drug treatment and behavioral assay. (B) Bar graph showing the discrimination index between a novel and familiar arm in the Y-maze test after 28 days of chronic ICV infusion of CXCL10 or CXCL10+AMG487. Data were plotted as the mean ± SEM, one-way ANOVA followed by Tukey’s multiple comparisons test, F _(2,12)_ =9.560, ***p<0.001 Veh vs. CXCL10; *p<0.05, CXCL10 vs. CXCL10+AMG487, n=6–8 mice/group. (C) A representative bar graph showing the discrimination index between novel mice and familiar mice after chronic treatment with CXCL10. Data were plotted as the mean ±SEM, unpaired t-test *p<0.05, t=2.390, Veh vs CXCL10, n= 6-7 mice/group. (D) Representative bar graph showing reduced percent freezing due to chronic treatment of mice with CXCL10, suggesting impaired learning and memory. Data were plotted as the mean ± SEM, one-way ANOVA followed by Tukey’s multiple comparisons test, F _(2,21)_ = 11.66, ***p< 0.001, Veh vs. CXCL10; **p<0.01, CXCL10 vs. CXCL10+AMG, n=6–8 mice/group. (E) Bar graph shows non-significant changes in locomotor activity due to chronic ICV infusion of CXCL10. Data were plotted as total distance traveled (cm)/60-minute and presented as mean ± SEM, n=8-9 mice/group. (F) Representative immunoblots showing increased expression of the senescence protein p16^INK4a^ in CXCL10-treated mice. (G) Bar graph showing quantification of p16^INK4a^ expression normalized to β-Actin, presented as percent vehicle (%Veh). Data were plotted as mean ± SEM, one-way ANOVA followed by Tukey’s multiple comparisons test, F _(2,13)_ =3.3340, *p<0.05, Veh vs. CXCL10, *p<0.05, CXCL10 vs. CXCL10+AMG, n=4-6 mice/group. (H) Representative immunoblots showing increased expression of the senescence protein p21^Cip1^ in CXCL10-treated mice. (I) Bar graph showing quantification of p21^Cip1^ normalized to β-Actin presented as percent vehicle (%Veh). Data were plotted as mean ± SEM, one-way ANOVA followed by Tukey’s multiple comparisons test, F _(2,21)_ =11.66, ***p<0.001, Veh vs. CXCL10; *p<0.05, CXCL10 vs. CXCL10+AMG, n=6-7 mice/group. (J) Representative immunoblots showing increased expression of p53 in the treated group as compared to Veh. (K) Bar graph showing quantification of p53 normalized to β-Actin presented as a percent vehicle (%Veh). Data were plotted as mean ± SEM, one-way ANOVA followed by Tukey’s multiple comparisons test, F _(2,14)_ =7.574, **p<0.01 Veh vs CXCL10, n= 5-7 mice/group. (L) Representative immunoblots showing decreased expression of pRB in CXCL10-treated mice. (M) Bar graph showing quantification of pRB normalized to β-Actin presented as percent vehicle (%Veh). Data were plotted as mean ± SEM, one-way ANOVA followed by Tukey’s multiple comparisons test F _(2,15)_ = 16.00, ****p<0.0001, Veh vs. CXCL10; *p<0.05 CXCL10 vs. CXCL10+AMG487, n=5-7 mice/group.

### 3. CXCL10 attenuates c-FOS expression in PFC

Immediate early genes (IEGs) such as c-FOS, Arc, and zif268 are rapidly induced within neurons after synaptic activation and have long been used as markers of behaviorally activated neurons (Herdegen and Leah, 1998; Katche et al., 2010). Multiple lines of evidence suggest that prefrontal neurons (such as those labeled by c-Fos) express high levels of c-FOS at the onset of learning and are crucial for the expression of memory (DeNardo et al., 2019; Kitamura et al., 2017). Given that increased CXCL10 in the brain inhibited learning and memory, we sought to determine if this alteration was due to reduced neuronal activation/synaptic plasticity. We used c-FOS promoter-driven eYFP expression (pAAV-c-FOS-eYFP) in the PFC **(Figure 4A)** as a surrogate marker for neuronal activation and network communications in mice. We observed that ICV infusion of CXCL10 for 28 days leads to significantly reduced expression of c-FOS-eYFP in the PFC, which was completely blocked by CXCR3 selective antagonist AMG487 **(Figure 4B, C)**.

**Figure 4:**
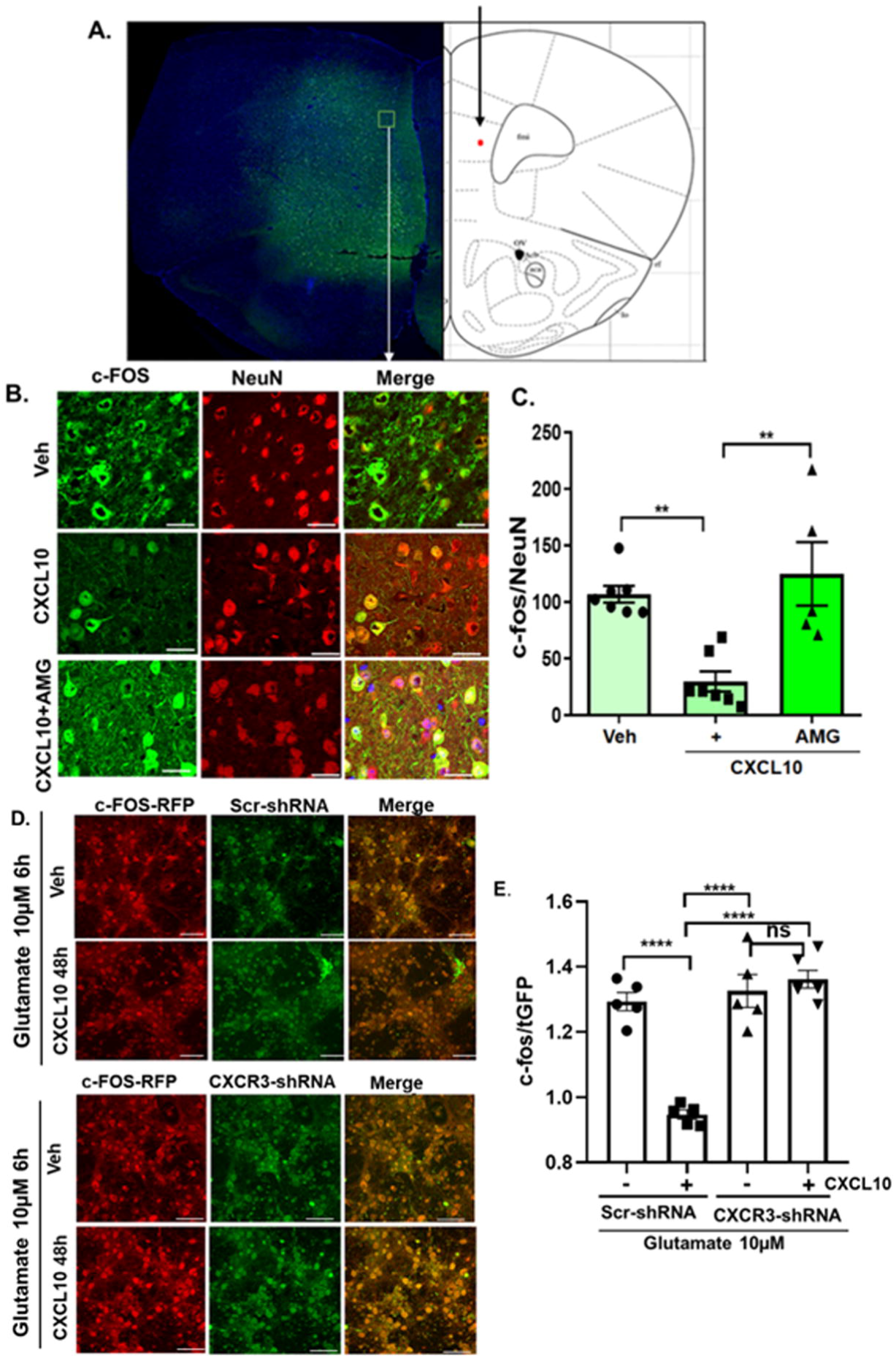
Chronic CXCL10 brain infusion impairs neuronal activation in PFC. (A) Representative immunohistochemistry images illustrating unilateral AAV-c-Fos-eYPF (green) stereotaxic injection in the mPFC of mouse brain, Anterior-posterior (AP) =+1.96mm, Dorso-ventral (DV)=-2.28mm, and Medial Lateral (ML)=±0.062mm. (B) Representative confocal Images (40X magnification) showing c-Fos-eYPF expression to represent spontaneous neuronal activity in CXCL10-treated mice (8-month-old C57BL/6J mice), c-Fos (green) co-localizing with the neuronal marker, NeuN (red), scale bar, 20µm. (C) Bar graph showing quantification of the relative fluorescent intensity of c-Fos-eYFP (green) normalized to NeuN (red) intensity. Data were plotted as the mean ± SEM, one-way ANOVA followed by Tukey’s multiple comparisons test, F _(2,16)_ =12.15, **p<0.001, Veh vs. CXCL10; **p<0.001, CXCL10 vs. CXCL10+AMG487, n=5-7 mice/group. (D) Representative confocal images (40X magnification) showing glutamate (10µM) induced expression of c-Fos-iRFP to represent neuronal activity in veh or CXCL10 (50ng/ml, 48hrs) pretreated primary cortical neurons. Upper panel neurons were transduced with scr-shRNA lentivirus, while lower panel were transduced with CXCR3-shRNA for 48hrs. (E) Bar graph showing quantification of the relative fluorescent intensity of c-Fos-iRFP (red) normalized to turbo-GFP in CXCR3-shRNA and Scrambled (Scr)-shRNA (green). Data were plotted as a mean ± SEM, one-way ANOVA followed by Tukey’s multiple comparisons test, F _(3, 17)_ = 35.50, ****p<0.0001, Scr-shRNA vs. Scr-shRNA+CXCL10 (48hrs), ****p< 0.0001, Scr-shRNA+CXCL10 (48hrs) vs. CXCR3-shRNA+CXCL10 (48hrs), ****p<0.0001, Scr-shRNA+CXCL10 (48hrs) vs. CXCR3-shRNA+CXCL10, (48hrs), n=5/group.

Furthermore, considering the intracerebral glutamatergic neurotransmission is crucial for higher-order cognitive functions and found to be altered in aging-associated disorders (Zott and Konnerth, 2023), we sought to determine the effect of CXCL10/CXCR3 signaling on glutamate-induced neuronal activation in primary cortical neurons using c-FOS promoter-driven iRFP (pAAV-c-FOS-iRFP) expression. Interestingly, we found that CXCL10 pre-treatment (50 ng/ml, 48hrs) significantly inhibited the glutamate (10 µM, 6hrs) induced c-FOS-iRFP expression, and surprisingly no effect of glutamate in CXCR3 knockdown neurons, suggesting the specificity of CXCL10 action through CXCR3 **(Figure 4D, E)**. To further determine if constitutive CXCR3 signaling plays a role in glutamatergic neurotransmission, we measured glutamate-induced (10 µM, 6hrs) c-FOS-iRFP) expression in control (Scr-shRNA transduced) and CXCR3 knockdown neurons. We found significantly increased expression of c-FOS-iRFP in CXCR3 knockdown neurons in comparison to scrambled shRNA transduced neurons **(Figure S4)**.

However, no difference in the extent of glutamate-induced c-FOS-iRFP expression was observed in control and CXCR3 knockdown neurons. These results clearly suggest that CXCL10/CXCR3 signaling attenuates glutamatergic neurotransmission in the brain.

### 4. CXCL10/CXCR3 signaling induces neuronal senescence and attenuated autophagy in primary neurons

To further elucidate if the CXCL10 acts specifically through CXCR3 and consequently induces senescence, we used primary cortical neurons 8-9 days *in vitro* (DIV) treated with either vehicle or CXCL10 (50 ng/ml). Our immunoblotting experiment revealed that CXCL10 treatment significantly induced p16^INK4a^ expression that was blocked by CXCR3 antagonist AMG487 (10 µM) **(Figure 5A-D)**. We also confirmed that CXCL10 induces the other markers of senescence, such as senescence-associated β-galactosidase (SA-β-gal) activity and lipid accumulation in neurons (Hamsanathan and Gurkar, 2022) **(Figure S6-7)**. Neurons rely heavily on autophagy to meet high metabolic demands, and decreased autophagy has been widely implicated in aging-associated CNS disorders, including AD dementia (Menzies et al., 2015). Considering these facts and our results of CXCL10-induced neuronal senescence, we also determined the effect of CXCL10/CXCR3 signaling on autophagy flux using primary cortical neurons prepared from CAG-RFP-EGFP-LC3 transgenic mice (RRID: IMSR_JAX:027139). As shown in **figure 5E-F**, CXCL10 treatment significantly increased green puncta due to decreased formation of autophagolysosomes and hence increased EGFP/RFP ratio, which was blocked by CXCR3 specific antagonist AMG487 (10 µM). Furthermore, the specificity of CXCL10/CXCR3 signaling on senescence induction was also confirmed in primary cortical neurons that were transduced either with AAV9-scrambled-shRNA or AAV9-CXCR3-shRNA for 72 hrs. As anticipated, CXCL10 treatment (50 ng/ml, 48 hrs) significantly induces p16^INK4a^ expression through CXCR3 receptors **(Figure 6)**.

**Figure 5:**
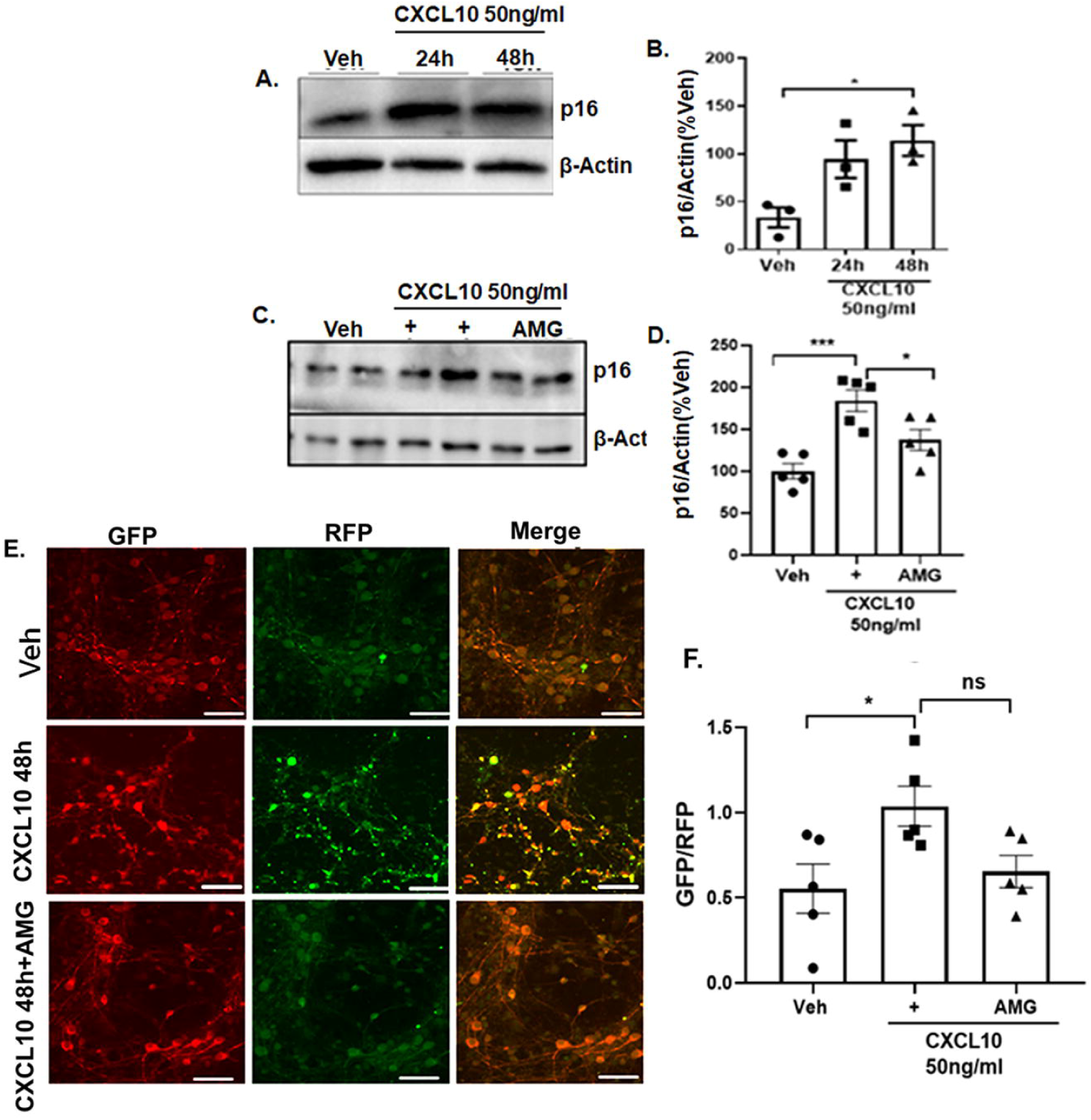
CXCR3 activation induces senescence and attenuates autophagy in primary cortical neurons. (A) A representative immunoblot shows a time-dependent increase in the expression of p16^INK4a^ due to CXCL10 (50 ng/ml) in primary cortical neurons. (B) Bar graph showing quantification of p16^INK4a^ normalized to β-Actin, presented as percent vehicle (%Veh). Data were plotted as mean ± SEM, one-way ANOVA followed by Tukey’s multiple comparisons test F _(2, 6)_ = 7.025, *p<0.05, Veh vs. CXCL10, (48hrs), n = 3/group (C) Representative immunoblot showing increased expression of senescence protein p16^INK4a^ due to CXCL10, which was inhibited by CXCR3 antagonist AMG487 (10 µM) for 48hrs. (D) Bar graph showing quantification of p16^INK4a^ normalized to β-Actin, presented as percent vehicle (%Veh). Data were plotted as mean ± SEM, one-way ANOVA followed by Tukey’s multiple comparisons test, F _(2,12)_ =13.28, ***p<0.001, Veh vs. CXCL10 (48hrs), *p<0.05, CXCL10 vs. CXCL10+AMG487(48hrs), n=5/group. (E) Representative confocal images (40X) of immunocytochemistry RFP (red) and GFP (green) showing LC-3 II expression in transgenic RFP-EGFP-LC3-tagged mice pups’ primary cortical neurons at 48hrs after treatment with CXCL10. (F) Bar graph showing quantification of GFP/RFP ratio for LC3-II. Data were plotted as mean ± SEM, one-way ANOVA followed by Tukey’s multiple comparisons test, F _(2,12)_ =4.474, *p<0.05, Veh vs. CXCL10 (48hrs), n=5/group.

**Figure 6:**
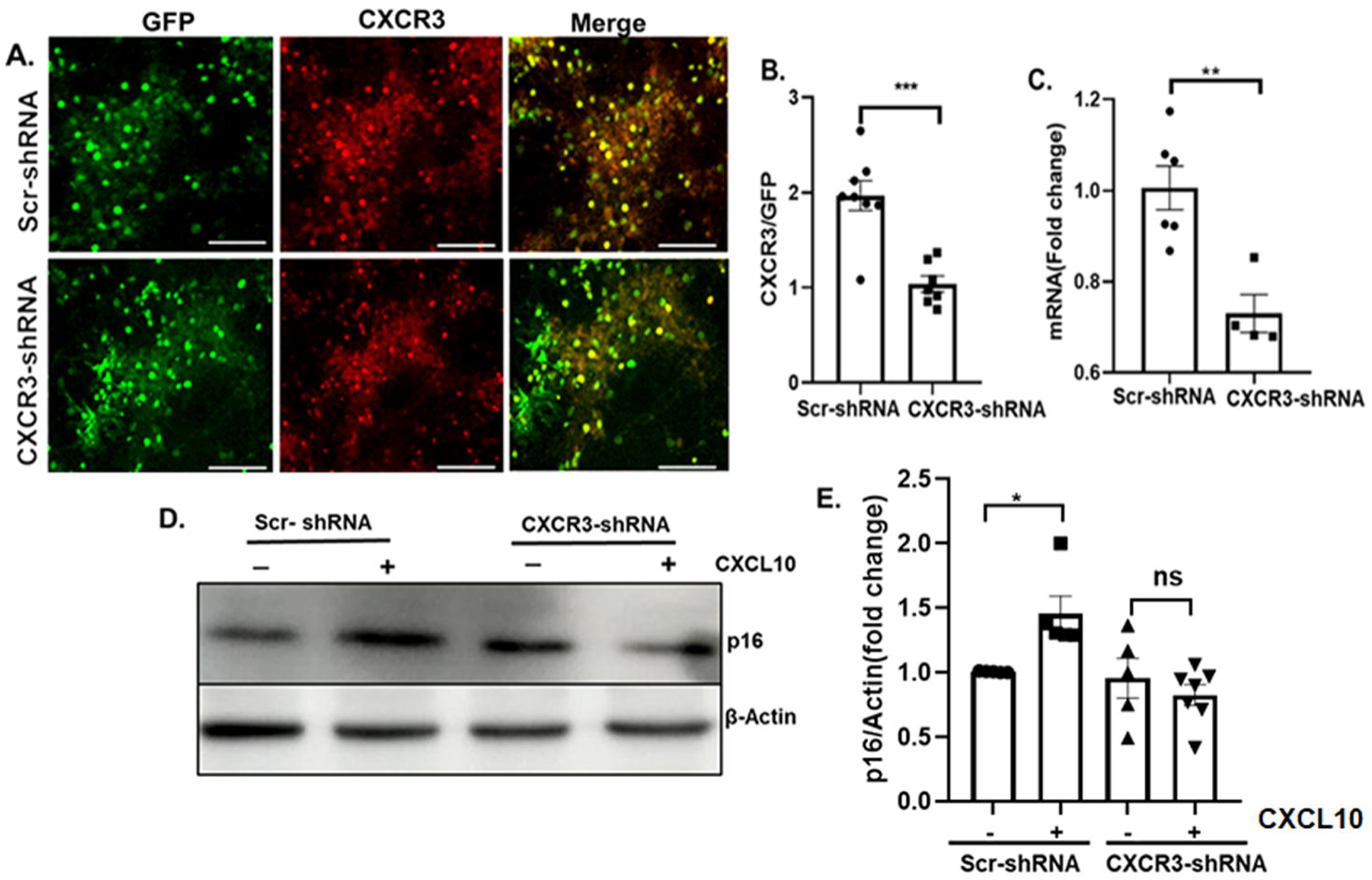
CXCR3 knockdown attenuates CXCL10 induced neuronal senescence. (A) Representative fluorescence (20X) images of immunocytochemistry, CXCR3 (red) and GFP (green), showing shRNA-mediated knockdown of CXCR3 in primary cortical neurons (B) Bar graph showing quantification of fluorescence intensity of turbo-GFP and CXCR3 (red) following shRNA-mediated knockdown of CXCR3 presented as the CXCR3/tGFP ratio. Data were plotted as a mean ± SEM, unpaired t-test, t=2.145 ***p<0.001, Scr-shRNA vs. CXCR3-shRNA, n=7-8/group. (C) Bar graph showing quantification of CXCR3 expression normalized to turbo-GFP presented as fold change following the lentivirus-mediated knockdown of CXCR3. Data were plotted as the mean ± SEM, unpaired t-test, t=3.180, **p<0.01, Scr-shRNA vs. CXCR3-shRNA, n=4-6/group. (D) Representative immunoblot showing the expression of p16^INK4a^ in primary cortical neurons treated with veh or CXCL10 (50 ng/ml) following Scr-shRNA or CXCR3-shRNA mediated transduction. (E) Bar graph showing quantification of p16^INK4a^ expression normalized to β-Actin presented as fold change. Data were plotted as mean ± SEM, one-way ANOVA followed by Tukey’s multiple comparisons test, F _(3, 13)_ =3.135, *p<0.05, Scr-shRNA vs. Scr-shRNA+CXCL10, (48hrs), n=5-7/group.

## Discussion

The seminal work of (Hayflick, 1965) for the first time showed a relationship between senescence and aging, and subsequent studies demonstrated that senescence itself can drive aging. Cellular senescence is now considered a fundamental aging mechanism that immensely contributes to aging-associated CNS disorders. Therapeutic strategies to either selectively eliminate the senescent cells or inhibit their secretome have lately gained much traction for neurodegenerative disorders (Chaib et al., 2022; Kennedy et al., 2014). In this study, using young (2M), middle-aged adult (12M) and old age (18M) C57BL/6J mice, we first established that increased levels of cellular senescence in PFC and hippocampus, critical brain regions involved in cognition. As expected, we also observed significant cognitive impairments in 18M-old mice. Interestingly, increased cellular senescence in both brain regions was found to be associated with increased levels of CXCL10 and its cognate receptor CXCR3 expression. To determine if there is any causal relationship between CXCL10 and cellular senescence/cognitive impairments, we administered a very modest amount of CXCL10 in the brain for 28 days (0.5pg/hrs) in 8M adult healthy mice and demonstrated that increased expression of cellular senescence marker proteins (p16^INK4a^, p21^Cip1^, p53) and cognitive impairment as we observed in 18M old mice. Importantly, the effect of CXCL10 on the expression of senescence markers and cognition was blocked by CXCR3 selective antagonist AMG487, clearly indicating a causative relationship between CXCL10/CXCR3 signaling and brain aging. Furthermore, using primary cortical neurons, we mechanistically established that increased CXCL10/CXCR3 signaling in neurons specifically induces senescence, decreased autophagy flux, and glutamatergic neurotransmission.

Unequivocal evidence from several studies strongly links chronic inflammation with cognitive impairments (Kipnis et al., 2004; Ziv et al., 2006). Also, studies with heterochronic parabiosis mouse models having increased circulating pro-inflammatory cytokines showed impaired cognition, which supports the concept of inflammaging (Villeda et al., 2011). Brain inflammaging is a chronic, low-grade inflammation that’s a key characteristic of age-related neurodegenerative changes. It is primarily characterized by the buildup of senescent cells and cognitive decline. Senescent cells secrete a variety of proteins, including pro-inflammatory cytokines and chemokines, such as CCL11, CXCL10, and IL-8 (Gluck et al., 2017; Villeda et al., 2011) (Acosta JC et al., 2018), into the surrounding fluid, which is known as the senescence-associated secretory phenotype (SASP). Although expression of both CXCL10 and its main receptor CXCR3 has been reported in neurons and glia (Krauthausen et al., 2015; Vinet et al., 2010), the role of CXCL10/CXCR3 signaling in inflammaging is not clear. Interestingly, higher circulating levels of CXCL10 in older cohorts have been reported to be associated with decreased DNA CpGs methylations within the CXCL10 gene promoter (Bradburn et al., 2018). Additionally, in this study, subjects with higher CXCL10 exhibited impaired spatial working memory, highlighting the important role of CXCL10 in aging-associated disorders.

However, the direct causative and mechanistic nature of the relationship between CXCL10 and brain aging / cognitive impairments is yet to be elucidated. In view of the above facts, our observation of increased expression of CXCL10, and CXCR3 in PFC and hippocampus, and impaired cognition in old age (18M) mice further validate the hypothesis of increased SASPs in brain inflammaging. Taking one step further, we determined if increased CXCL10/CXCR3 signaling alone is capable of inducing cellular senescence in the healthy brain, and indeed we found that modest brain infusion of CXCL10 induces senescence markers (p16^INK4a^, p21^Cip1^, p53, decreased pRB) in CXCR3 dependent manner. Furthermore, we also observed a significant decrease in c-FOS driven eYFP reporter in PFC, a hallmark of neuronal activation, suggesting a dampened neuronal network activity, which is in contrast to earlier reports of in AD patients and acute brain inflammation (Busche et al., 2015; Kastanenka et al., 2020). This discrepancy could stem from the selective inhibition of CXCL10/CXCR3 signaling in cortical neurons as opposed to amyloid plaques causes both glial and neuronal perturbations in AD.

Brain senescence has been usually attributed to the senescence of astrocytes and microglia, which ultimately interfere with glial–neuron communication (Crowe et al., 2016; Safaiyan et al., 2016). However, later, it was also found that post-mitotic neurons also follow the same senescence pathways, i.e., p53/p21^Cip1^ and p16^INK4a^/pRB tumor suppressor pathways (Jurk et al., 2012; Simpson et al., 2010; Simpson et al., 2016; Vazquez-Villasenor et al., 2021). In this context, we also observed that increased CXCL10/CXCR3 signaling induces primary cortical neuron senescence via both p53/p21^Cip1^ and p16^INK4a^/pRB tumor suppressor pathways.

Although we didn’t determine the effect of CXCL10/CXCR3 signaling on glial functions and consequently affects neuronal functions, it is quite likely that some of the deleterious effects of increased CXCL10 signaling is contributed by microglia (Krauthausen et al., 2015). Glutamate, an important excitatory neurotransmitter of the CNS, acts mostly in the hippocampus and cortex and is known to play an important role in learning and memory formation (Kinoshita et al., 2016; Zott and Konnerth, 2023). Although decreased glutamatergic neurotransmission in subcortical regions due to inflammation is reported to cause decreased psychomotor speed, reaction time, and information processing, and ultimately, depression-like conditions (Haroon et al., 2016; Haroon et al., 2017). However, it is not clear how inflammaging affects glutamatergic signaling in the brain. Therefore, we sought to determine the effect of CXCL10/CXCR3 signaling on glutamatergic signaling in primary cortical neurons using a c-FOS-iRFP reporter. Our results clearly demonstrated that selective activation of CXCL10/CXCR3 signaling suppresses glutamate-induced neuronal activation, suggesting a significant inhibitory effect of inflammaging on glutamatergic neurotransmission. Although we have not investigated in this study, the inhibitory effect of chronically increased CXCL10/CXCR3 signaling could be due to decreased levels of Brain-Derived Neurotrophic Factor, which supports glutamatergic synaptic transmission and plasticity. In conclusion, this study reveals a novel role of CXCL10 in the brain, which is the induction of inflammaging.

## Acknowledgments

All Authors are thankful to the Director of CSIR-CDRI, for providing the necessary research facilities. M.P. acknowledges the AcSIR and CSIR, New Delhi, for research fellowships. The writers acknowledge Mr. Devanshu Kaushik and Mrs. Deepmala Umrao’s editing assistance. Additionally, the authors acknowledge the Sophisticated Analytical Instrumentation Facility & Research (SAIF & R) for supplying useful Intravital Imaging Facility Olympus BX61-FV1200-MPE data, and the in-house Animal House Facility for behavioral investigations. Addgene provided the created plasmids pAAV-c-FOS-iRFP and pAAV-c-Fos-eYFP, for which the authors are also grateful. The CSIR-CDRI communication number for this manuscript is-161/2024/PNY

## Author Contributions

Monika Patel-Manuscript writing, study design, and conducting the experiment for figures 1-6, Sakesh Kumar-ICV cannula and Alzet pump installation (Figure. 3). Aditya Singh-Characterization of transgenic (RFP-EGFP-LC3) mice (Figure. 5), Prem Narayan Yadav-Design of experiments, data interpretation, and manuscript writing.

## Abbreviations

ANOVA-Analysis of variance, ACSF-Artificial Cerebrospinal Fluid, DMSO-Dimethyl sulfoxide, PFA-Paraformaldehyde, AD-Alzheimer’s Disease-CNS, Central Nervous system, CXCL10-, C-X-C motif chemokine ligand 10, CXCR3, C-X-C motif chemokine receptor 3, SASPs-Senescence associated secretory phenotypes, PFC-prefrontal cortex.

