## supplementary data for "CXCL10/CXCR3 Signaling Induces Neural Senescence and Cognitive Impairments"

**Table S1.** List of primers used for RT-PCR

| **S.N.** | **Primer name** | **Sequence (5’ - 3’)** |
| --- | --- | --- |
| **Primers used for RT-PCR** | | |
| 1 | tGFP-F | TACTACAGCTCCGTGG |
| 2 | tGFP-R | TTCTTCACCGGCATCTGCAT |
| 3 | CXCR3-F | GGATCTATTTCCGGTGAATTCGCCACCATGAAGACGATCA |
| 4 | CXCR3-R | CTAGAACTAGTCTCGAGGAATTCCTAGAGGCCAGAATAACTAGCC |

**Table S2.** List of antibodies and its dilution used in various experiments

| **Antibody Name** | **Dilutions** | **Source** | **Catalog No.** | **Application** |
| --- | --- | --- | --- | --- |
| Anti-p16^INK4a^ | 1:1000 | Invitrogen | PAZ-20379 | WB/ICC |
| Anti-p21^Cip1^ | 1:1000 | Invitrogen | AHZ0422 | WB |
| Anti-LC3 | 1:1000 | Sigma | L7543 | WB |
| Anti- p53 (Invitrogen, cat. no. MA5-12571, 1:1000) | 1:1000 | Invitrogen | MA5-1257 | WB |
| Anti-pRB | 1:1000 | Cell Signaling Technology  TTechnology | 9307 | WB |
| Anti-CXCR3 | 1:250 | Abcam | ab71864 | IHC/ICC |
| Anti-CXCL10 | 1:250 | Abcam | ab9938 | IHC |
| Anti-NeuN | 1:500 | Cell Signaling Technology | 94403 | IHC/ICC |
| Anti-NeuN | 1:500 | Abcam | ab128886 | IHC |
| Anti-RFP | 1:200 | Abs | AB-2802 | ICC |
| Anti-tGFP | 1:100 | Origine | TL500382V | ICC |
| Anti-MAP2 | 1:1000 | Sigma | M9942 | ICC |
| Anti-c fos | 1:500 | Abcam | ab7963 | IHC |

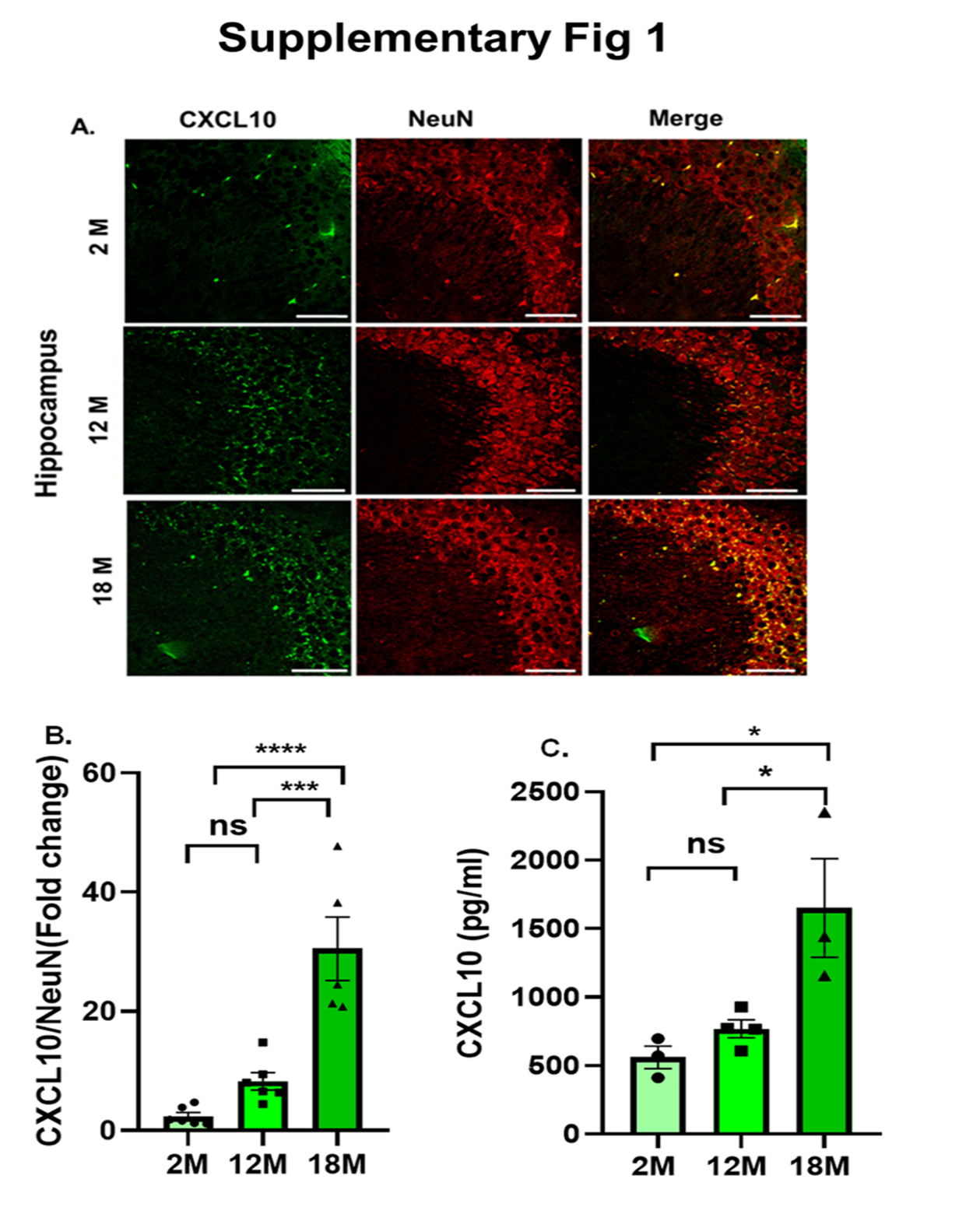

**Figure S1: Increased CXCL10 expression in an age-dependent manner in the hippocampus (A)** Representative confocal images (40X magnification) of the hippocampus showing increased CXCL10 expression in 12M and 18M old C57BL/6J mice, CXCL10 (green) co-localizing with the neuronal marker NeuN (red), scale bar, 20µm. **(B)** Bar graph showing quantification of relative fluorescent intensity of CXCL10, normalized to NeuN intensity. Data were plotted as the mean ±SEM, one-way ANOVA followed by Tukey's multiple comparisons test F _(2, 14)_ = 24.06, ***p<0.001, 2M vs. 18M; **p<0.01, 12M vs. 18M, n=5-6 mice/group. **(C)** A significant increase in the level of CXCL10 in the hippocampus was also observed by the ELISA test. Data were plotted as ±SEM, one*-*way ANOVA followed by Tukey’s multiple comparisons test, F _(2, 7)_ =8.265, *p<0.05, 2M vs. 18M, *p<0.05, 12M vs. 18M, n=3-4 mice/group.

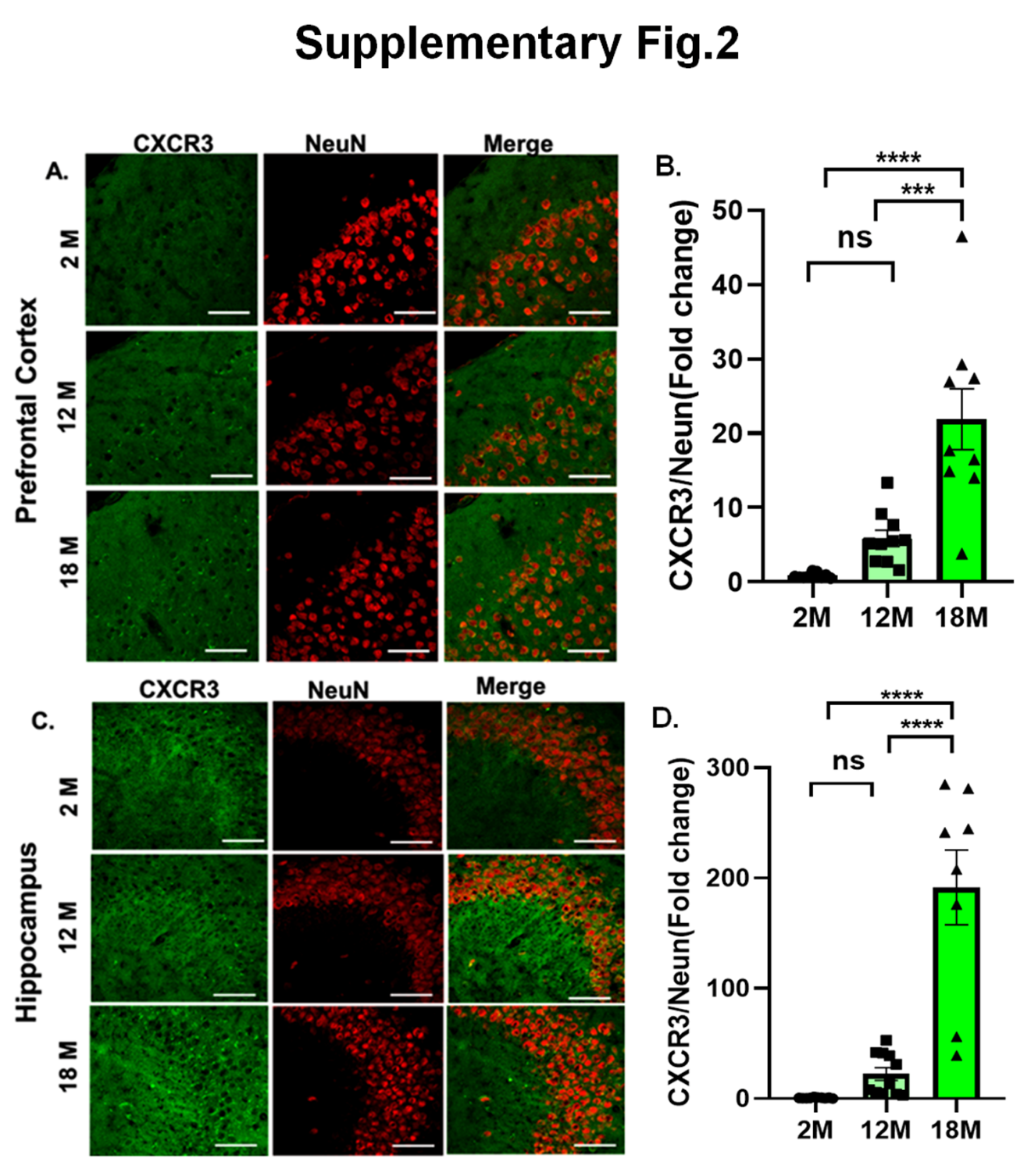

**Figure S2: Increased CXCR3 expression in PFC and hippocampus.** (A**) Representative** confocal images (40X magnification) of the PFC showing increased expression of CXCR3 in 12M and 18M old-C57BL/6J mice, CXCR3 (green) co-localizing with the neuronal marker NeuN (red); scale bar 20µm. **(B)** Bar graph showing quantification of relative fluorescent intensity of CXCR3, normalized to NeuN intensity. Data were plotted as mean ± SEM, one-way ANOVA followed by Tukey's multiple comparisons test, F _(2, 25)_ = 20.8, ****p<0.0001(2M vs. 18M), ***p<0.001 (12M vs.18M), 8-9 mice/group. **(C)** Representative confocal images (40X magnification) of the hippocampus showing increased expression of CXCR3 in 12M and 18M old- C57BL/6J mice, CXCR3 (green) co-localizing with neuronal marker NeuN (red), scale bar, 20µm. **(D)** Bar graph showing quantification of relative fluorescent intensity of CXCR3, normalized to NeuN intensity. Data were plotted as mean ± SEM, one-way ANOVA followed by Tukey's multiple comparisons test, F _(2, 25)_ = 35.60, ****p<0.0001 (2M vs. 18M), ****p<0.0001 (12M vs. 18M), 8-9 mice/group per.

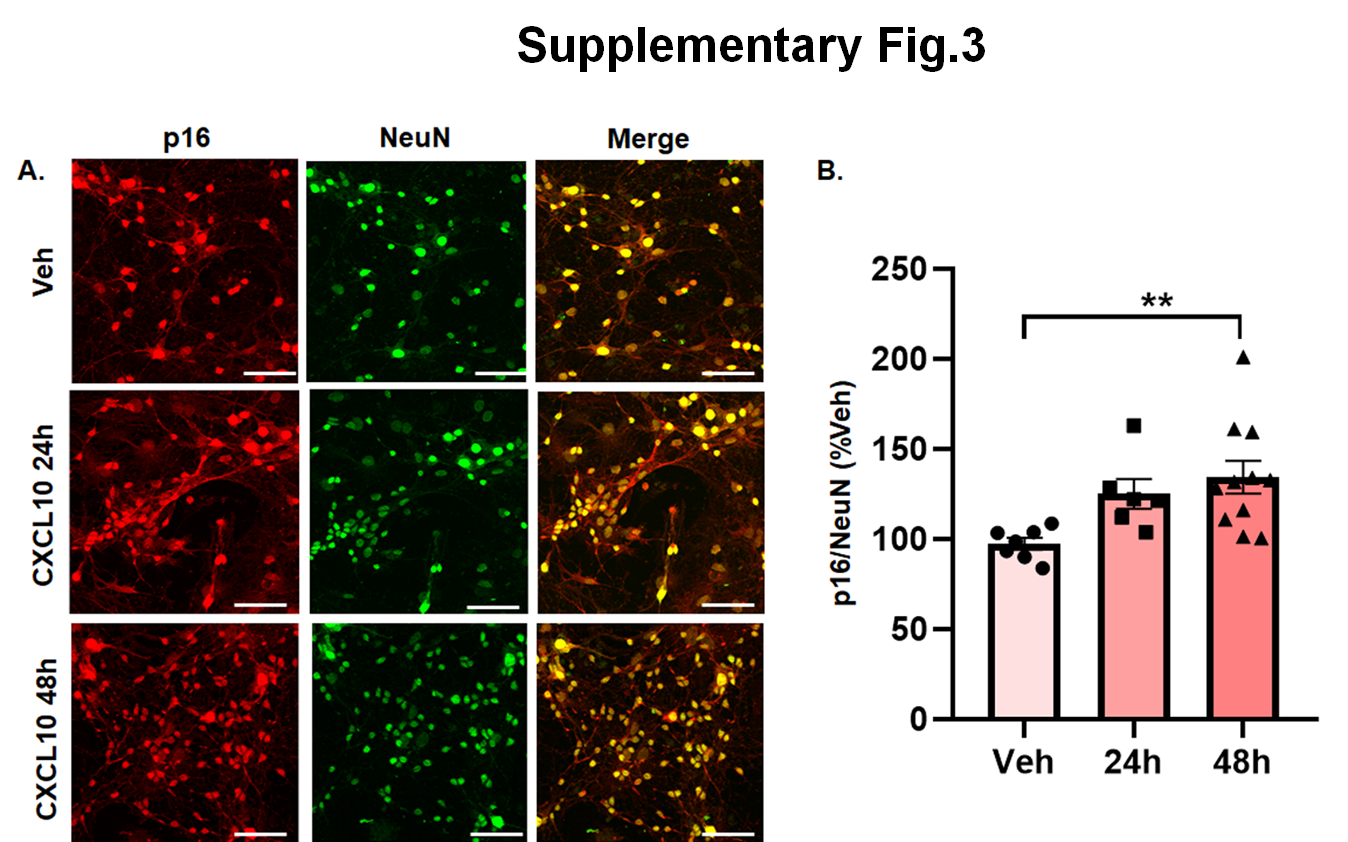

**Figure S3: CXCR3 activation induces senescence marker p16^INK4a^ in primary cortical neuron. (A)** Representative confocal images (40X magnification) of immunocytochemistry showing increased expression of p16^INK4a^ in primary cortical neurons treated with CXCL10 (50 ng/ml) at 48hrs, p16^INK4a^ (red) co-localizing with the neuronal marker NeuN (red); scale bar, 20 µm. **(B)** Bar graph showing quantification of relative fluorescent intensity of p16^INK4a^, normalized to NeuN intensity. Data were plotted as mean ± SEM, one-way ANOVA followed by Tukey’s multiple comparisons test, F _(2, 21)_ = 5.428, **p<0.01, Veh vs. CXCL10 (48hrs), n =7-11/group.

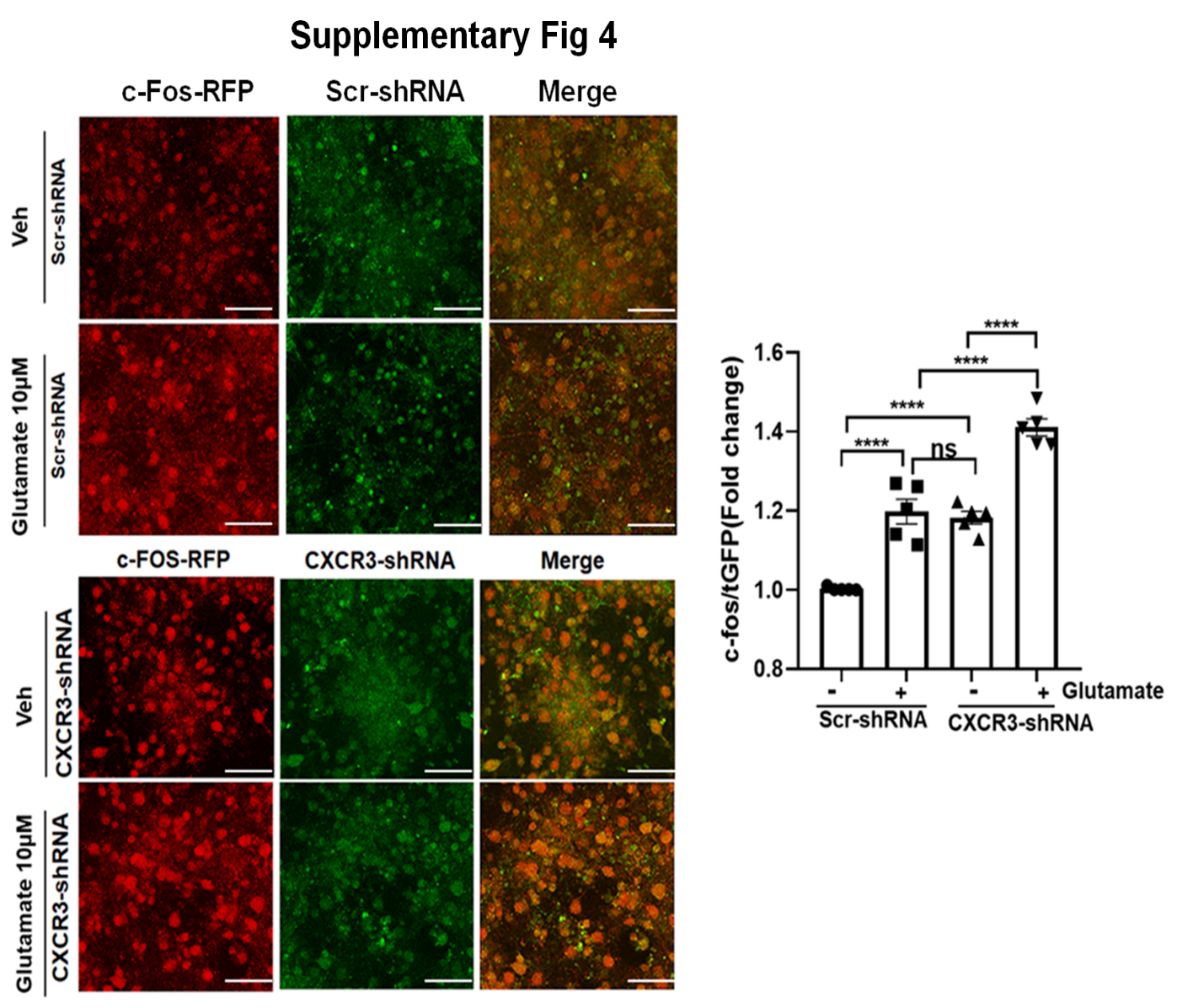

**Figure S4: CXCR3 knockdown accelerates the expression of c-Fos in primary cortical neurons. (A)** Representative confocal images (40X magnification) showing glutamate (10µM) induced expression of c-Fos-iRFP to represent neuronal activity in veh or CXCL10 (50ng/ml, 48hrs) pretreated primary cortical neurons. Upper panel neurons were transduced with scr-shRNA lentivirus, while lower panel were transduced with CXCR3-shRNA for 48hrs. **(B)** Bar graph showing quantification of the relative fluorescent intensity of c-Fos-iRFP (red) normalized to tGFP in CXCR3-shRNA and Scrambled (Scr)-shRNA (green). Data were plotted as mean ± SEM, one-way ANOVA followed by Tukey’s multiple comparisons test, F _(3, 19)_ = 97.42, ****p<0.0001, n=5/group.

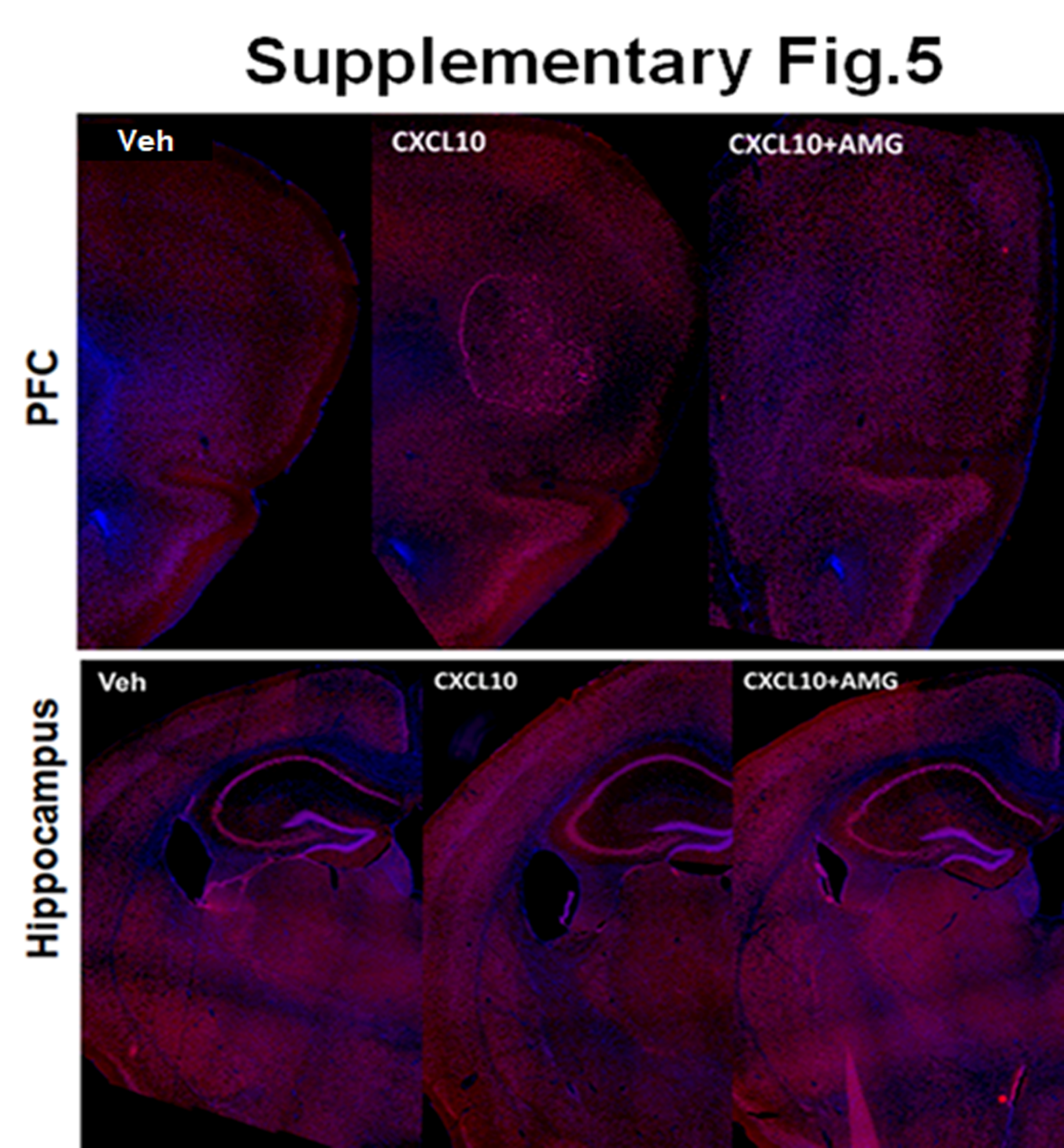

**Figure S5: CXCL10 infusion didn’t induce gross anatomical changes.** Representative PFC and hippocampal coronal sections of mice brain section were stained with Hoechst 33258 (Blue) and NeuN (Red) and fluorescent montage images were captured using 10X objectives Leica-DMI6000.

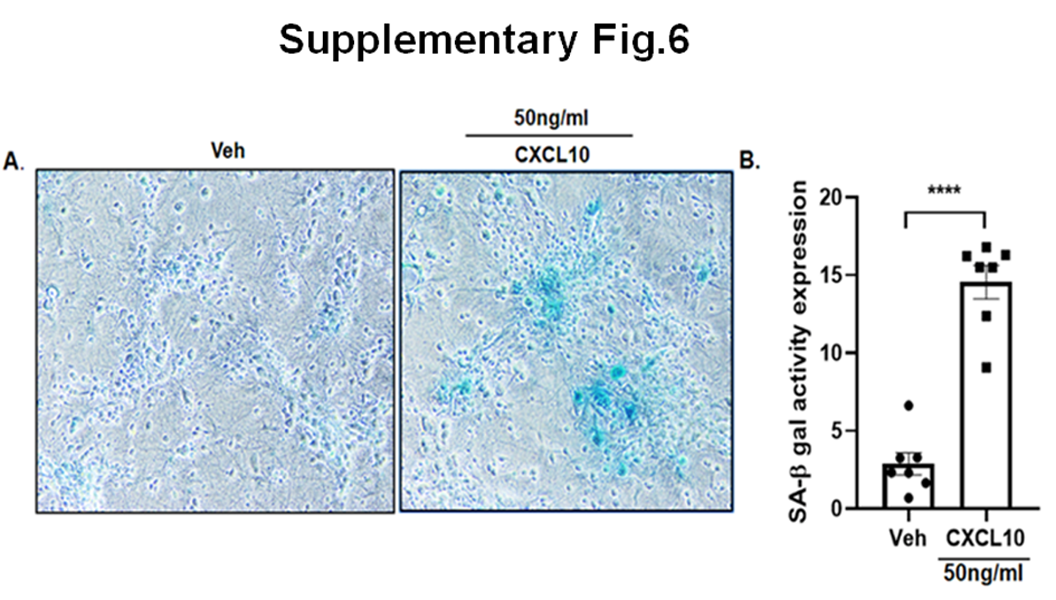

**Figure S6: CXCL10 induced SA-ꞵ gal activity in primary cortical neuron** (A) Representative 20X bright field images showing expression of SA-ꞵ gal staining (blue color) in primary cortical neuron after treatment with veh or CXCL10 for 48hrs. (B) Bar graph showing the quantification of SA-ꞵ gal staining (blue color). Data were plotted as unpaired t- test, t=9.120, ****p<0.0001, Veh vs CXCL10 (48hrs), n=6-7/group.

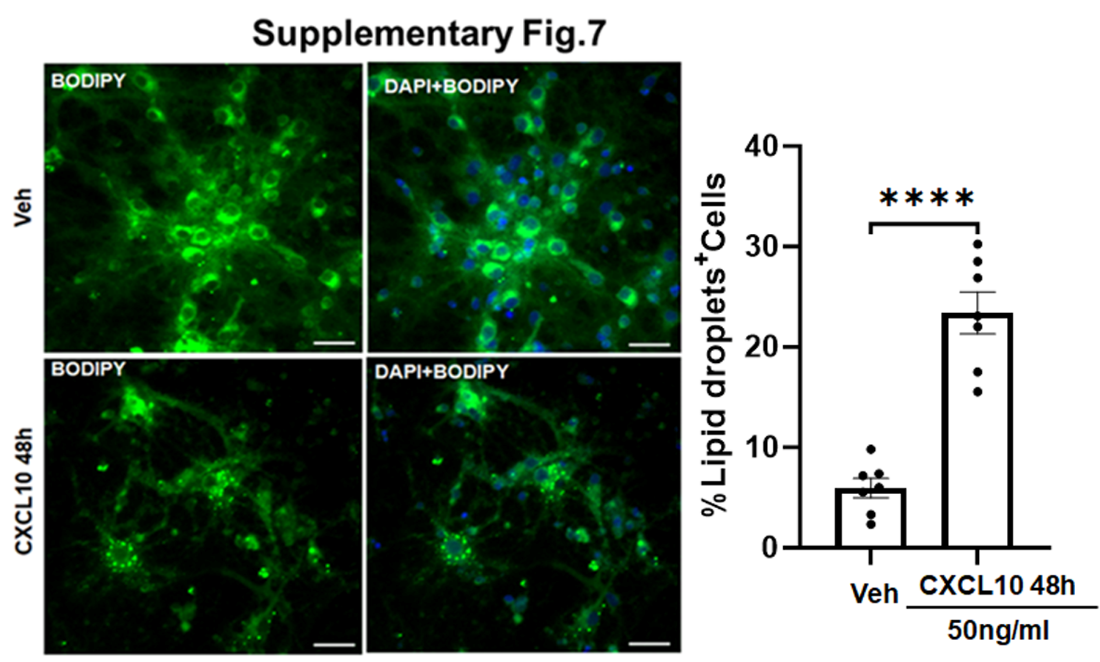

**Figure S7: CXCL10 induced lipid droplets accumulation in primary cortical neuron** (A) Representative 20X fluorescence images showing expression of lipid droplets accumulation (green color) in primary cortical neuron after treatment with veh or CXCL10 for 48hrs. (B) Bar graph showing the quantification of lipid droplets positive cells (green color). Data were plotted as %lipid droplet^+^ cells in veh or CXCL10 treated cells, ****p<0.0001(Veh vs CXCL10), unpaired t- test, t=7.590, n=7/group.

**Supplementary Materials and Methods**

**1.1 Immunohistochemistry**

C57BL/6J mice were perfused with a solution of 4% paraformaldehyde through their hearts. After removal, their brains were fixed overnight in 4% paraformaldehyde and then dehydrated in 30% sucrose for 24 hours. Thin coronal slices of 20 μm were obtained using a cryostat (FSE, Thermo Scientific). To stain the slices, they were washed four times in PBS and subjected to antigen retrieval in 10mM sodium citrate buffer (pH-6.3), followed by permeabilization using 0.5% Triton X-100 in 1X PBS for one hour at room temperature. After that, the slices were blocked with a buffer consisting of 3% bovine serum albumin, 3% horse serum, and 0.3% Triton X-100 in PBS for two hours at room temperature. Finally, the slices were incubated overnight at 4°C with anti-rabbit CXCR3 (Abcam, ab71864) with dilution 1:500, and anti-mouse NeuN (CST, 94403) with 1:500. After an overnight incubation, the brain sections were washed three times with PBST (0.1% Triton X-100 in 1X PBS). Then, the sections were incubated with respective secondary antibodies that were conjugated with Alexa fluor 594 or 488 (1:1000) and Hoechst 33258 (2µg/ml) in a blocking buffer for 1 hour at room temperature. After incubation, the slices were washed thrice, and they were then mounted onto Poly-L-lysine-coated glass slides using a Vectashield antifade mounting medium. Fluorescent images were collected with XYZ acquisition mode at 1μm step size using the Olympus BX61-FV1200-MPE microscope, which used a 40X oil (1.3 NA) objective at 8.0μs/pixel scan speed (Olympus, Shinjuku, Tokyo, Japan). To quantify the pixel intensity for CXCR3 (Abcam, ab71864) and NeuN (CST, 94403) staining Hippocampus, pixel intensity quantification was performed using ImageJ software (www.rcb.info.nih.gov/ij). The results were presented as fold change after normalization with pixel intensity of NeuN in each region.

**1.2 ELISA**

We used the CXCL10 ELISA kit from R&D Systems (DY466-05) for measuring the levels of mouse CXCL10 following the manufacturer’s instructions. Firstly, we homogenized mouse hippocampal tissue lysates in lysis buffer -150 mM NaCl, 50 mM Tris, 1%Triton X-100, with protease inhibitors. Then, we centrifuged the homogenized mixture at 10000g for 20 minutes at 4°C and collected the supernatants, which were stored at -80°C until further use. Next, we coated a 96-well microplate with the diluted capture antibody according to the kit instructions and sealed the plate for an overnight incubation at room temperature. The following day, the microplate was washed three times with wash buffer (0.05% Tween 20 in PBS, pH 7.2) and blocked with blocking buffer (1% BSA in PBS pH 7.2) for one hour at room temperature. Thereafter, we incubated the plate with 100 µg of tissue lysates for 2 hours at room temperature. We then washed the microplate twice with wash buffer and added the detection antibody for 2 hours. We repeated the washing steps and incubated the plate with conjugated streptavidin-HRP for 30 minutes at room temperature, followed by washing the plate and adding substrate solution for 20 minutes at room temperature. Lastly, we added a stop solution to each well and read the plate with a microplate reader at 450 nm. The data was analyzed by flex station 3 a molecular device.

**1.3 Immunocytochemistry and confocal imaging for primary cortical neuron**

Primary cortical neurons were prepared from 0–1-day-old pups and the cells were fixed with 2% PFA for 10 minutes at 12 days *in vitro* (DIV). To stain the culture, we washed them four times in PBS, and permeabilized them using 0.3% Triton X-100 in 1X PBS for an hour at room temperature. Next, we blocked the primary cortical neuron using a buffer containing 3% bovine serum albumin, 3% horse serum, and 0.3% Triton X-100 in PBS for two hours at room temperature. We incubated primary culture overnight at 4°C using at a dilution of 1:500, anti-mouse NeuN (CST, 94403) at a dilution of 1:200, and anti-rabbit p16^INK4a^ (Invitrogen, PAZ-20379) at a dilution of 1:500. After overnight incubation, primary cortical neurons were washed three times with PBST (0.1% Triton X-100 in 1X PBS). After that, primary culture was mixed with the appropriate secondary antibodies that were linked to Alexa fluor 594 or 488 (1:1000) and Hoechst 33258 (2 µg/ml) in a blocking buffer. This was done at room temperature for one hour. Following the incubation, we washed the slices three times before mounting them onto poly-lysine-coated glass slides using a vectashield antifade mounting medium. Fluorescent images were collected with XYZ acquisition mode at 1μm step size using the Olympus BX61-FV1200-MPE microscope, which used a 40X oil (1.3 NA) objective at 8.0 μs/pixel scan speed (Olympus, Shinjuku, Tokyo, Japan). To quantify the pixel intensity for p16^INK4a^ and NeuN staining in primary cortical neurons, pixel intensity quantification was performed using ImageJ software (www.rcb.info.nih.gov/ij). The results were presented as a percent Vehicle after normalization with the pixel intensity of NeuN and p16^INK4a^.

**1.4 Senescence associated β-galactosidase activity**

SA-β-gal staining was performed as described earlier by Dimiri et al 1995. Primary cortical neurons treated with vehicle or CXCL10 for 48hrs were fixed with 4% paraformaldehyde in PBS (pH 7.2) for 15 min at 4°C followed by 3 washes with PBS. Thereafter, cells were incubated in 0.1m-phosphate buffer (pH 6.0) containing 5mM potassium ferrocyanide, 5mM potassium ferricyanide, 150 mM NaCl, 2 mM MgCl2, 1mg/ml (X-gal, SRL-45904) (5-bromo-4-chloro-3-indolyl-ꞵ-D-galactopyranoside) substrate for 24 hours at 37°C without CO2 in darkness. After incubation, cells were again washed with PBS buffer and mounted with Vectashield antifade mounting media. The image was acquired using Nykon Eclips TS100 light field microscopy using 20X objectives, and the blue color intensity was quantitated using ImageJ software (www.rcb.info.nih.gov/ij).

**1.5 Lipid droplets staining in primary cortical neuron**

Primary cortical neuronal cells were seeded at 1×10^6^ cells on poly-L-lysine-coated glass coverslips in complete neurobasal media. Following specific treatments, cells were fixed in 4% PFA for 30 min, washed 3x in PBS and incubated in PBS with BODIPY™ 493/503 (1:1000 from 1mg/ml stock solution in DMSO; Thermo Fisher) and Hoechst 33258 (2 µg/ml) (1:1000) for 15 min at RT. A total of three randomly selected visual fields per coverslip were photographed (20x magnification) using a fluorescence microscope (Leica DMI600). To analyze the percent lipid droplets containing primary cortical neurons, the total number of Hoechst+ cells with lipid droplets were counted in 7 images (from a total of 3 independent experiments) by an experimentally blind observer, and data were presented as percent vehicle (%veh).
